# Differential encoding of temporal context and expectation under representational drift across hierarchically connected areas

**DOI:** 10.1101/2023.06.02.543483

**Authors:** David G Wyrick, Nicholas Cain, Rylan S. Larsen, Jérôme Lecoq, Matthew Valley, Ruweida Ahmed, Jessica Bowlus, Gabriella Boyer, Shiella Caldejon, Linzy Casal, Maggie Chvilicek, Maxwell DePartee, Peter A Groblewski, Cindy Huang, Katelyn Johnson, India Kato, Josh Larkin, Eric Lee, Elizabeth Liang, Jennifer Luviano, Kyla Mace, Chelsea Nayan, Thuyanhn Nguyen, Melissa Reding, Sam Seid, Joshua Sevigny, Michelle Stoecklin, Ali Williford, Hannah Choi, Marina Garrett, Luca Mazzucato

## Abstract

The classic view that neural populations in sensory cortices preferentially encode responses to incoming stimuli has been strongly challenged by recent experimental studies. Despite the fact that a large fraction of variance of visual responses in rodents can be attributed to behavioral state and movements, trial-history, and salience, the effects of contextual modulations and expectations on sensory-evoked responses in visual and association areas remain elusive. Here, we present a comprehensive experimental and theoretical study showing that hierarchically connected visual and association areas differentially encode the temporal context and expectation of naturalistic visual stimuli, consistent with the theory of hierarchical predictive coding. We measured neural responses to expected and unexpected sequences of natural scenes in the primary visual cortex (V1), the posterior medial higher order visual area (PM), and retrosplenial cortex (RSP) using 2-photon imaging in behaving mice collected through the Allen Institute Mindscope’s OpenScope program. We found that information about image identity in neural population activity depended on the temporal context of transitions preceding each scene, and decreased along the hierarchy. Furthermore, our analyses revealed that the conjunctive encoding of temporal context and image identity was modulated by expectations of sequential events. In V1 and PM, we found enhanced and specific responses to unexpected oddball images, signaling stimulus-specific expectation violation. In contrast, in RSP the population response to oddball presentation recapitulated the missing expected image rather than the oddball image. These differential responses along the hierarchy are consistent with classic theories of hierarchical predictive coding whereby higher areas encode predictions and lower areas encode deviations from expectation. We further found evidence for drift in visual responses on the timescale of minutes. Although activity drift was present in all areas, population responses in V1 and PM, but not in RSP, maintained stable encoding of visual information and representational geometry. Instead we found that RSP drift was independent of stimulus information, suggesting a role in generating an internal model of the environment in the temporal domain. Overall, our results establish temporal context and expectation as substantial encoding dimensions in the visual cortex subject to fast representational drift and suggest that hierarchically connected areas instantiate a predictive coding mechanism.

## 1 Introduction

Neural populations in the visual cortical hierarchy encode specific features of visual stimuli, such as orientation, spatial frequency, and direction of movement [Hubel and Wiesel, 1959, 1962, Siegle et al., 2019, de Vries et al., 2020]. Recent studies have also shown more diverse encoding capacities of the visual cortex. For example, activity in visual cortex exhibits strong modulation by changes in behavioral state [Niell and Stryker, 2010, Stringer et al., 2018, Musall et al., 2019, Salkoff et al., 2020, Nestvogel and McCormick, 2022, Poort et al., 2015], arousal [McGinley et al., 2015], and attention [Ito and Gilbert, 1999, Thiele et al., 2009, True, 2001, McAdams and Reid, 2005, Poort et al., 2021]. Trial and reward history have also been shown to influence visual responses [McMahon and Olson, 2007, Meyer et al., 2014, Nikolić et al., 2009, Shuler and Bear, 2006, Ramadan et al., 2022, Gillon et al., 2021], indicating that sensory coding is influenced not only by current state but also prior experience and expectations. Neurons in primary visual cortex (V1) learn short spatiotemporal sequences of stimuli upon repeated presentation [Gavornik and Bear, 2014], and enhance their activity for unexpected oddball images, as well as at the start of a novel sequence [Homann et al., 2022, Kim et al., 2019b].

Predictive coding has been proposed as a theory that can account for contextual modulation of sensory responses [Khan et al., 2018, Keller and Mrsic-Flogel, 2018], framing sensory perception as a process of active inference. The predictive coding framework proposes that connections between hierarchically organized areas operate to construct a model of the environment by comparing bottom-up sensory inputs with top-down prior experience and expectations to continually update the representation of the environment [Rao and Ballard, 1999, Bastos et al., 2012, Friston, 2005]. Accordingly, areas providing feedback to sensory regions should represent learned expectations, while early sensory areas should encode deviations from these expectations.

Retrosplenial cortex is an association area providing feedback input to the visual cortex [Harris et al., 2019, Van Groen and Wyss, 2003, Wyss and Van Groen, 1992], and in turn receives input from visual cortex [Sit and Goard, 2022, Murakami et al., 2015, Van Groen and Wyss, 2003] as well as the hippocampus and other medial temporal areas [Sugar et al., 2011, Wyss and Van Groen, 1992], serving as a bridge between sensory and cognitive representations. The retrosplenial cortex is involved in memory, spatial navigation, and prospective thinking [Vann et al., 2009, Alexander and Nitz, 2015]. It was recently proposed that the diverse functions ascribed to retrosplenial cortex could be unified under a theory such as the predictive coding framework, based on its ability to generate predictions through integration of sequences of stimuli experienced in time [Alexander et al., 2022]. In support of this idea, [Makino and Komiyama, 2015] showed that top-down projections from retrosplenial cortex to V1 are strengthened with experience, and develop a ramping profile predictive of a learned event.

One potential challenge to studying the mechanisms of predictive coding is the presence of representational drift, a gradual change in neural representations which has been found in multiple brain regions [Schoonover et al., 2021b, Driscoll et al., 2017], including V1 [Deitch et al., 2021, Aitken et al., 2022]. Predictive coding relies on the accuracy of bottom up sensory representations, such that internal models can be updated when the environment deviates from expectations. Accordingly, instability in representations may interfere with accurate updating. However, the presence of drift is not incompatible with predictive coding, and instead may be critically related. Representational drift has been proposed as a substrate for continual learning [Aitken et al., 2022, Rule et al., 2019, Driscoll et al., 2022], which is critical to updating in predictive coding. Even in primary visual cortex, drift in representations may reflect meaningful contextual differences between repeated presentations of the same stimulus that need to be incorporated into internal models, or be reflective of plasticity in other parts of the network [Rule et al., 2020].

The effects of temporal context, expectation, and drift have been investigated so far in different experimental setups. Here, we perform a comprehensive experimental and computational analysis uncovering the interaction between temporal context, expectation and representational drift on visual responses along three hierarchically connected areas: the primary visual cortex (V1), the posterior medial higher order visual area (PM), and retrosplenial cortex (RSP). Area PM sits between V1 and RSP and is highly interconnected with both areas [Wang et al., 2012, Harris et al., 2019, Van Groen and Wyss, 2003], raising the possibility that PM relays information about natural scene statistics to RSP and information about learned expectations back to V1, and may display intermediate characteristics between the two. We recorded neural activity in these areas as mice viewed repeated natural image sequences, image sequence violations, and image pairs outside the repeated sequence order. We found that visual responses encoded three main effects: temporal context (recent scene transition history), expectation (sequence violation), and representational drift. The features exhibited by such contextual effects were strongly area-specific and revealed complex interactions, consistent with the theory of hierarchial predictive coding.

First, we found that natural image encoding depended on the history of transitions preceding each image (temporal context) and cortical regions. Decodability of images identities decreased along the cortical hierarchy. Specifically, neurons in V1 and PM could clearly encode image identity in any temporal contexts, while in RSP, decodability of image identity decreased significantly. Generally, within the same region, information about image identity was enhanced when images were presented in pairs or in longer sequences, compared to when they were presented in randomized order in V1 and PM. The contribution of temporal context to encoding of image identity, however, varied across cortical regions.

Second, we found that visual responses were strongly affected by expectations. In the main sequence block, a decoder trained on visual responses in V1 and PM could robustly distinguish expected from unexpected images when the expected image was replaced with an unexpected oddball one, and oddball responses were strongly enhanced compared to expected ones. However, population responses to oddballs in RSP recapitulated the expected image that was replaced. These results are consistent with predictive coding, with sensory areas (V1 and PM) signaling deviations from expectation, and higher order areas (RSP) recapitulating predictions based on expectations and on past history.

Third, we found a strong drift of population activity on the timescale of minutes within the recording session, whose features differed along the hierarchy. In V1 and PM, we found that drift, defined by a significant encoding of elapsed time within the session, also preserved the representational geometry of the evoked responses, consistent with previous experimental and modeling studies [Deitch et al., 2021, Aitken et al., 2021, Qin et al., 2023]. On the other hand, RSP showed significant encoding of elapsed time within a session, but the representational geometry of evoked responses to sensory stimuli did not generalize across epochs, suggesting that relevant dimensions for RSP activity may not be stimulus specific, but potentially more related to overall environmental context. Importantly, we found that drift could not be explained away by behavioral measures in any of the three areas.

Together, these results provide evidence that temporal context and expectation are differentially represented across hierarchically connected areas in a manner consistent with the predictive coding framework, with V1 and PM encoding image identities and transitions as well as image sequence violations and RSP encoding expected stimuli. The hierarchical features of representational drift were also consistent with predictive coding, maintaining stable encoding of bottom-up sensory representations. The complex pattern of coding we observed across V1, PM, and RSP, provides a more complete picture of the interactions between temporal context, expectation, and representational drift at different levels of a hierarchically connected circuit.

## 2 Results

We set out to investigate how neural responses to natural images depend on temporal context and expectation, and whether the effects of temporal context differ across levels of an interconnected hierarchy. We defined temporal context as the set of transitions preceding each presented image, representing the short-term history of stimulus presentation. We designed a stimulus protocol in which four natural images were presented in different temporal contexts (250ms stimulus with no interleaving gray screen): either in random order (‘randomized control’), or in a four image main sequence (ABCD, denoted ‘sequence’ hereafter), or in randomized pairs of images, recapitulating the transitions between images in the main sequence (‘transition control,’ with AB, BC, CD, DA, CX*_i_*, X*_i_*A pairs randomly interleaved, for *i* = 1*, …,* 10 oddballs). In the “sequence” block, ten rare, “oddball” images randomly replaced the fourth image of the set to form an unexpected sequence (ABCX; Fig. 1b). In contrast to previous studies investigating the effects of stimulus history on responses in visual cortex [Kim et al., 2019a], we used natural images instead of gabor patches. Each session featured four blocks, where randomized control occurred both as the first and the last block, and sequence and transition control blocks as second and third, respectively. Experiments using this stimulus protocol were conducted through the OpenScope program at the Allen Institute using a standardized pipeline for in vivo 2-photon calcium imaging (Fig. 1a).

**Figure 1.**
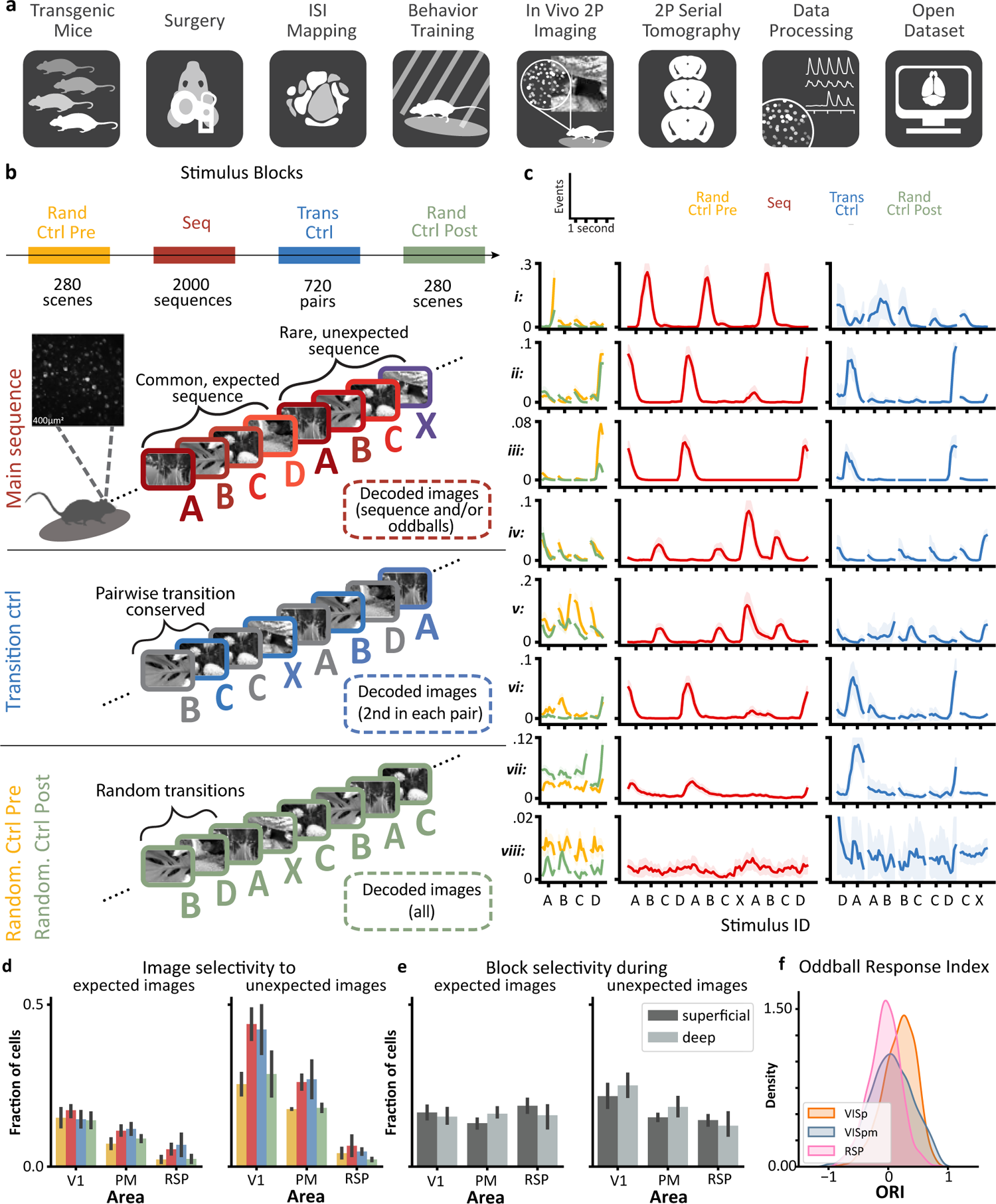
Presenting images in a variety of stimulus blocks determines single cell activity along the visual hierarchy. **a)** Experimental pipeline from developing transgenic mice, to recording population responses in the visual cortex, to finally an open dataset available to the public in the DANDI archive.**b)** Mice passively view natural images in different stimulus blocks while neural activity is recorded from V1, PM, or RSP. Images are either presented in random order (yellow and green) or in common, expected sequences with the occasional oddball interleaved (red), or in transition control (blue) which only preserves pairwise transitions. **c)** Single cell PSTHs to natural images in each stimulus blocks. Each row shows a representative average response from each recorded area, with the shaded area showing the variance across trials. i-ii: Superficial V1 neurons. iii-iv: Deep V1 neurons. v: Superficial PM neuron. vi: Deep PM neuron. vii: Superficial RSP neuron. viii: Deep RSP neuron. **d)** One-way ANCOVA results, accounting for the locomotion of the animal. Left: A small fraction of cells in V1, PM, and RSP are selective to expected images in all stimulus blocks. Right: A larger fraction of cells in V1 and PM are selective to unexpected images in the sequence and transition control blocks as compared to the randomized control ones. Error bars indicate standard deviation across depths. **e)** Using responses to expected images (ABCD) and unexpected images (X), we can assess whether a cell is selective to the stimulus blocks in which the image was presented. Error bars indicate standard deviation across different images. **f)** Histogram of Oddball Response Indices calculated for each cell in the 3 areas.

### 2.1 Mixed selectivity in single-cell responses

Populations of excitatory neurons were measured across multiple depths (range: 125 - 450 *µ*m), with a total of 2299 neurons in V1, 2071 neurons in PM, and 1628 neurons in RSP. We defined the ‘stimulus block’ as one of the four blocks within each session, presented in the following order: randomized ctrl pre, main sequence, transition ctrl, randomized ctrl post (Fig. 1b). Single cell responses across all areas and conditions were highly heterogeneous (Fig. 1c and d). Interestingly, we found hints of temporal context dependence and expectation in evoked responses already at the single cell level, suggesting complex mixed selectivity to multiple aspects of the experiment. In V1 and PM, we found cells that responded strongly to the presentation of a preferred ‘main sequence’ image (Fig. 1c rows i-vii) and/or to the unexpected ‘oddball’ image X*_i_* (for *i* = 1*, …,* 10; Fig. 1c rows iv,v). Whereas in RSP we found some cells that were selective to main sequence images, but not oddball ones (row vii), as well as cells that were not selective to any image but were selective to the block in which the image was presented in. This block selectivity extended to all areas for both during expected and unexpected images (Fig. 1e). Overall, a small fraction of cells were selective to expected images in all stimulus blocks (Fig. 1d). Moreover, in V1 and PM but not in RSP, larger fractions of cells were selective to unexpected images and responses to unexpected images were stronger than those to expected images (Fig. 1f), consistent with a bottom-up prediction error in the hierarchical predictive coding theory. Neurons in RSP were the least selective to expected and unexpected images among the areas. Interestingly, we found a large fraction of neurons across all 3 areas that were selective to the block in which the image was presented in (Fig. 1e). To understand how the interactions between expectation and temporal context shape visual representations, we further performed a series of population analyses based on cross-validated classification, described in the sections below.

### 2.2 Natural scene encoding varies with the transition history along the cortical hierarchy

First we set out to investigate how natural scenes are encoded across the visual hierarchy. We constructed cross-validated linear classifiers to decode image identity using population responses within each stimulus block (randomized control pre and post, transition control, main sequence). We used a linear classifier as it can be interpreted as representing the activity of a downstream neuron receiving projections from the observed populations; although all our results were confirmed using nonlinear classifiers. Crucially, here and in all subsequent decoding analyses, when comparing classification accuracy between conditions, we always matched the sample sizes of the respective training sets. Furthermore, we controlled for an animal’s behavioral state by comparing resting and running epochs.

We first focused on decoding the expected main sequence images within each stimulus blocks (ABCD; Fig. 2a). We found significant decoding of natural image identity during all of the stimulus blocks (sequence, transition control, and randomized control blocks) for all depths of V1 (Fig. 2b,d) and PM (Fig. 2d). However, in RSP, only supragranular (Fig. 2c) layers, but not infragranular layers significantly encoded the identity of expected images during the sequence and transition control stimulus blocks (Fig. 2d). During the randomized control blocks, we were not able to decode expected image identity in RSP across all depths. We show the full encoding accuracies across all stimulus blocks in both superficial and deep layers of V1, PM, and RSP in (Fig. S2).

**Figure 2.**
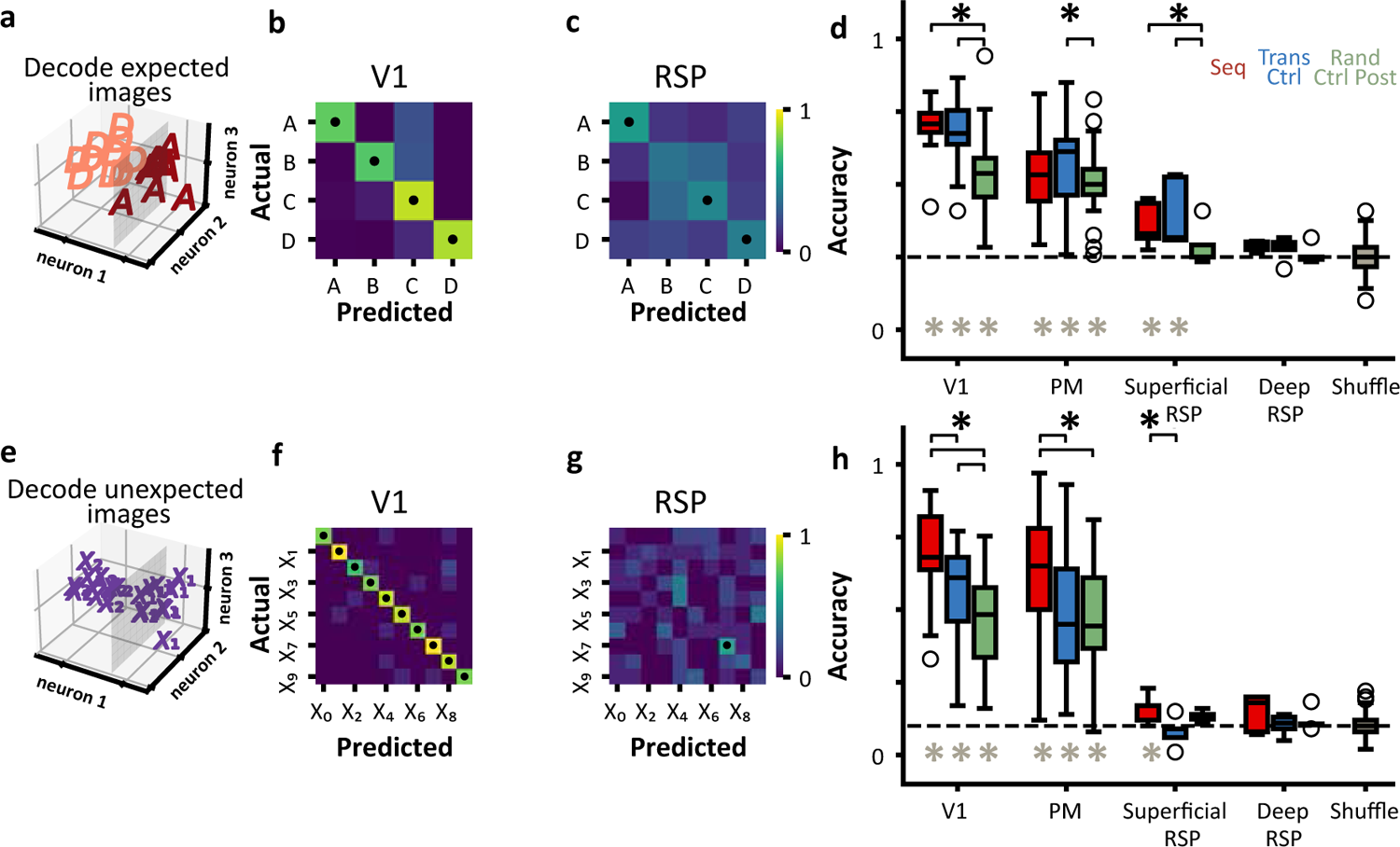
Decoding of natural images across different stimulus blocks. **a)** Schematic showing linearly separable population responses to natural images presented in the main-sequence. **b & c)** Example confusion matrices show significant decoding of main-sequence images using superficial V1 or RSP population responses during the sequence stimulus block. **d)** Significant decoding, denoted by the gray asterisk (t-test compared to shuffle distribution, p < 0.05), of expected images along the visual hierarchy in different stimulus blocks. All depths of V1 and superficial RSP, show an increase in decoding performance for the sequence and transition control blocks relative to randomized control (black asterisk, Wilcoxon signed-rank test, p < 0.05). In PM, only the transition control block was significantly different compared to randomized control. Deep RSP remains at chance level for all stimulus blocks. Randomized control pre not shown for clarity. **e)** Schematic showing linearly separable population responses to unexpected natural images. **f & g)** Example confusion matrices show significant decoding of unexpected oddball images using V1 population responses during the sequence stimulus block, but not for RSP population responses. **h)** V1 and PM show significant decoding of unexpected natural images in all stimulus blocks (gray asterisk). V1 shows an increase of decoding performance when unexpected images disrupt 2-image sequences and even more so to 4 image sequences, as compared to randomized control. In PM, decoding performance of unexpected images is larger for the sequence block compared to transition control and randomized control, which are statistically insignificant from each other. Superficial RSP shows slight decoding performance to oddball images in the sequence block, but no others. Deep RSP remains at chance level.

Overall, we found that the decoding performance in all of the stimulus blocks decreased along the visual processing hierarchy (Fig. 2d). While decoding of natural scenes was modulated by behavior, this general trend was preserved (Fig. S3). A series of control analysis on publicly available datasets from V1 and PM [de Vries et al., 2020, Siegle et al., 2021] further confirmed the significant encoding of natural images in both V1 and PM with decreased accuracy in PM. However, we also observed an overall decrease in image encoding accuracy in datasets obtained by calcium imaging compared to eletrophysiological recordings, suggesting limitations of the calcium imaging technique compared to electrophysiology (Fig. S1).

Interestingly, even within the same region, decoding accuracy of image identities varied significantly with ‘temporal context’ immediately preceding the presented image, defined by the *transitions* between consecutive images. Generally, image encoding was more pronounced in the main sequence and transition blocks, compared to the randomized control blocks. In V1 and superficial RSP, both the sequence and transition blocks showed significantly higher image decodability than the randomized controls, while in PM, only the transition control produced significantly higher decodability compared to the randomized controls. In infragranular layers of RSP, the encoding accuracy was very low for all stimulus blocks. Note that in the transition control block, the transition preceding each decoded image is fixed and image-specific (i.e., A always precedes B, B always precedes C, and so on; Fig. 1b). In the main sequence block, on the other hand, not only the transition immediately preceding the decoded image, but all transitions were fixed and image-specific. Therefore, the comparable encoding accuracy levels during the transition control and sequence blocks across all regions, along with the significant coding differences between transition control and randomized control, suggests that temporal context immediately preceding the presented image has a higher impact on encoding of natural scenes than extended temporal contexts (more than one image transition preceding the presented image). However, the significantly accurate identification of natural scenes in the randomized control block in V1 and PM emphasizes that the identity of present images is clearly encoded in V1 and PM responses without the temporal context of the preceding images, which only strengthens the present image representations in the neural activity.

Next, we asked whether encoding of natural image identity differed between expected and unexpected (oddball) images. We used cross-validated linear classifiers to decode oddball image identity from population responses within each stimulus block (Fig. 2e). We found that V1 (Fig. 2f) and PM, but not RSP (Fig. 2g), displayed significant encoding of oddball image identity in all stimulus blocks, with similar decoding accuracies compared to the decoding of main sequence image identity ABCD. Unlike in the case for the expected images, for oddball images, the decodability was not significantly higher than the chance level even in superficial RSP (Fig. 2h).

Contrary to decoding expected main sequence images ABCD from each other, where the temporal context of each decoded image (i.e., all preceding transitions) is fixed and image-specific, all oddball images X*_i_* were preceded by the same image C, thus eliminating the image-specificity of the transitions. Therefore, the relative improvement (Fig. 2d) in decoding accuracy of images ABCD from randomized control to transition control may be ascribed to the fixed image-specific transitions preceding each image. On the other hand, encoding of the oddball image identity cannot depend on the image-specific transitions preceding oddballs, since all oddballs were preceded by the same image C. Indeed, in V1 and PM, we observed that oddball identity can be decoded significantly better in the sequence blocks compared not only to the randomized control, but also to the transition control block (Fig. 2h), consistent with the increased single-cell selectivities observed in V1 and PM in (Fig. 1d). We thus hypothesized that the increased decodability of oddball identities in the lower cortical areas V1 and PM during the sequence block may be due to their unexpected presentations in a learned sequences. We explore the potential relation of image encoding to disruptions in expectation in section 2.3 and in Figure 3.

**Figure 3.**
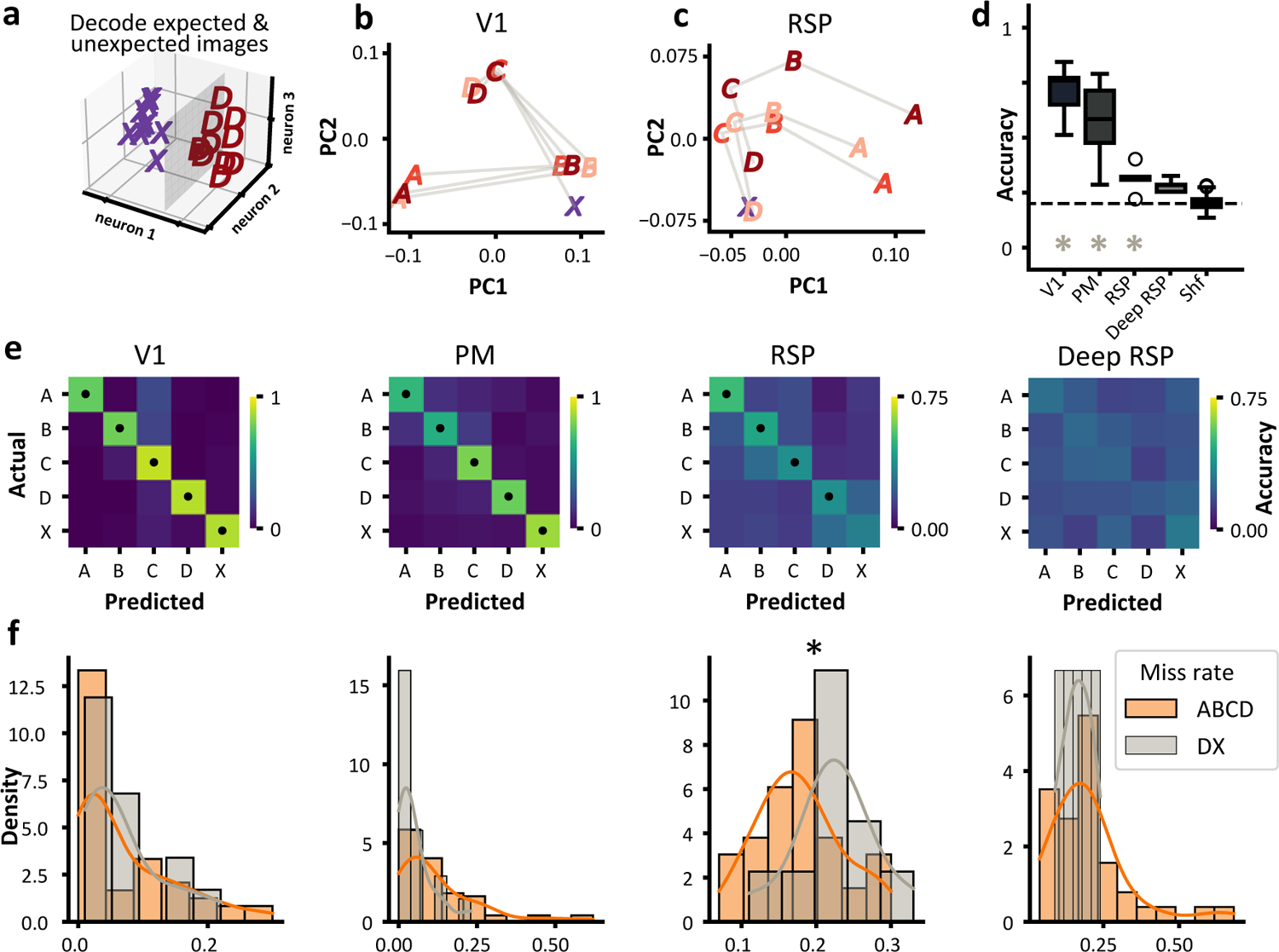
Decoding responses to expected and unexpected natural images reveals possible predictive coding mechanism in RSP. **a)** Schematic showing the task of classifier. **b)** PCA of V1 PSTHs for the sequences immediately preceding and following an unexpected event (X*_i_*). Oddball representation in PCA is distinct from expected image D. **c)** PCA of superficial RSP PSTHs for the sequences immediately preceding and following an unexpected event (X). Oddball representation in PCA space overlaps that of the expected image D. **d)** Significant decoding of main-sequence and oddball images along the visual hierarchy in the sequence. Hash pattern for RSP denotes deep layers (> 350 *µ*m) **e)** Example confusion matrices from superficial V1, superficial PM, and superficial/deep RSP. **f)** Histograms per area of the relative miss rates of main sequence images (A instead of B, B instead of C, etc) compared to that misclassifying images D and X. Superficial RSP reveals a significant increase in the DX miss-rate relative to the ABCD miss-rate.

### 2.3 Unexpected events disrupt encoding of natural scenes in RSP

In V1 and PM, single-cell image selectivity was larger for oddball responses X*_i_* compared to main sequence images ABCD (Fig. 1d), suggesting the presence of a surprise signal; on the other hand, RSP cells did not exhibit significant selectivity for oddball identity. We performed a series of population classification analysis to further characterize responses to unexpected images across areas and stimulus blocks (Fig. 2e-h). In V1 and PM, we found significant encoding of oddball images in all of the stimulus blocks (Fig. 2f). Stronger encoding during the sequence block compared to transition control shows that the temporal context of transitions preceding the decoded image enhance image encoding for unexpected images (Fig. 2h). On the contrary, we found that information about oddball image identity was not present in RSP population responses (Fig. 2g). Comparing to the small but significant decoding of expected images ABCD in RSP during the sequence block (Fig. 2c-d), the lack of information about unexpected oddball image identity suggests that expectations might affect RSP responses in a complex and different way, compared to V1 and PM.

We hypothesized that the decrease in unexpected image decoding compared to expected images in RSP could be due to predictive coding effects, whereas during the presentation of the unexpected sequence ABCX, the RSP population evokes a representation of the missing expected image D, in place of the unexpected X*_i_*. We first compared the representational geometry of the population responses to expected ABCD sequences vs. unexpected ABCX sequences across areas. In V1, we found that population responses encoded a unique representation of the oddball X, distinct from that of the missing expected image D (Fig. 3b; similar representations exist in PM, not shown). However, in RSP, the oddball representation was entirely overlapping with that of the missing image D (Fig 3c), suggesting that RSP might encode for the missing image D when presented the oddball during the sequence block.

To investigate the origin of this difference within the visual processing hierarchy, we simultaneously decoded both main sequence and oddball images within the sequence stimulus block. We hypothesized that, if oddball X evoked a representation of the missing D, the false positive rate of a decoder trained to classify X*_i_* from main sequence images ABCD would be larger for D than for the other ABC images, and indistinguishable from the D and X hit rates Jezzini et al. [2013]. We constructed a classifier using population responses to images that directly preceded an unexpected oddball image (..ABC*DABCX*ABCD…; Fig. 3a). The qualitative picture from the PCA analysis (Fig. 3b-c) was confirmed by our decoding results. Population responses in V1 and PM were able to discriminate between expected and unexpected images (Fig. 3d, e, f, V1 and PM). In superficial RSP, we found that the false positive rate of misclassifying oddball X*_i_* as the missing D was larger than misclassification with ABC (Fig. 3e, f, RSP). We performed a more granular pairwise analysis where we tested binary classification of each expected image (A, B, C, or D) vs oddballs X, and found that, while responses to images A, B, and C could be discriminated from responses to the oddballs, responses to image D was indistinguishable from the oddballs in RSP (Fig. S4). We concluded that in RSP, the unexpected oddball image X*_i_* evoked a representation that is indistinguishable from the missing image D. These results are consistent with an interpretation whereby RSP instantiates a predictive coding mechanism by evoking information about expected, but missing, visual signals (Fig 3c).

### 2.4 Population activity encodes information about stimulus blocks

The differential encoding of natural images across stimulus blocks naturally led us to ask whether there is contextual information on top of the representation of natural image identity or transition identity. We first tested whether single-cell responses could discriminate whether the same image was presented in different blocks (‘randomized control pre’,‘main sequence’,‘transition control’,‘randomized control post’). We found that single cells in all areas exhibited very pronounced selectivity for stimulus blocks, comparable to the selectivity for main sequence image identity (Fig. 1d and e). Remarkably, whereas RSP single cell responses were not selective for image identity (Fig. 1d), they strongly encoded the stimulus block each image was presented in (Fig. 1e).

To further quantify this, cross-validated linear classifiers were constructed based on population responses to the same image in each of six *epochs* that were constructed from the four stimulus blocks by splitting the sequence block into three epochs (‘early seq’, ‘middle seq’, ‘late seq’) (Fig. 4a). Using population responses to each of the main-sequence and oddball images, we were able to discriminate the stimulus block in which an image was presented in. This contextual information was present across all layers and depths recorded in V1 and PM (Fig. 4b) and was encoded on the comparable level for main sequence (expected) vs oddball (unexpected) images (Fig. S5a-d). In RSP, the contextual information is present in superficial layers but not in deep layers (Fig. 4b).

**Figure 4.**
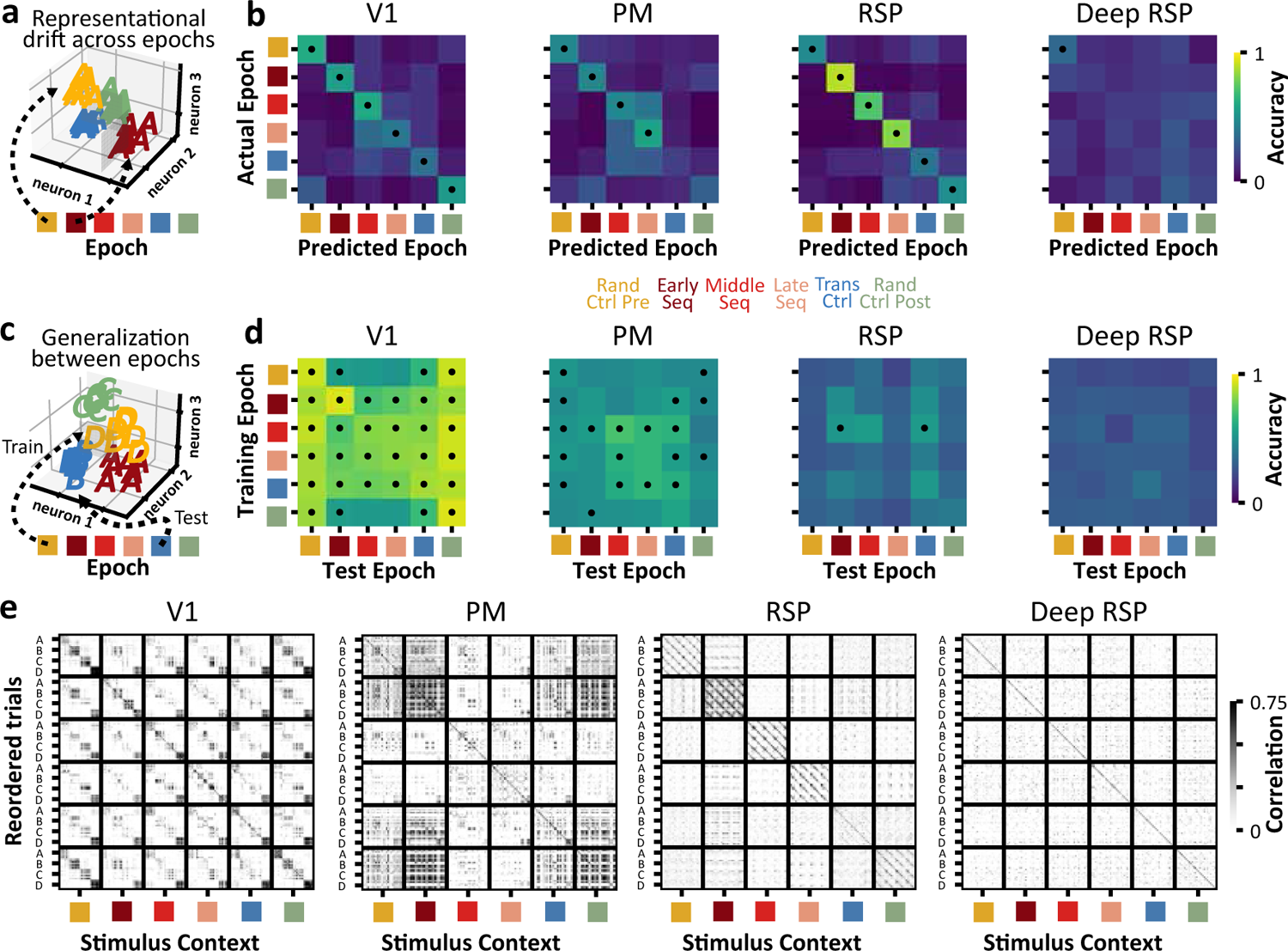
Generalization performance under representational drift. **a**) Schematic showing linearly separable population responses to the same natural image presented in different stimulus blocks. Decoding the time or stimulus block in which the same image is presented is our measure of representational drift. **b**) Example confusion matrices showing significant decoding of stimulus block using population responses from V1, PM, and superficial RSP. Representations from Deep RSP are unable to perform the decoding task. Epochs include the four main stimulus blocks, plus 2 additional epochs from early and late in the sequence block. **c**) Schematic showing population responses to main sequence images in different epochs of the session, with training and testing occurring using data from different stimulus blocks (epochs). **d**) Matrices show the generalization performance between epochs of the same sessions as panel *b*. Each element in the matrix represents a diagonal of a confusion matrix. In V1, representations of main sequence images generalize across stimulus blocks, while in PM, this phenomenon is less persistent. For superfical and deep RSP, generalization does not manifest. **e**) Example representational similarity matrices for V1, PM, and RSP show the correlation structure within and between stimulus blocks. Trials of main-sequence images are reordered to show correlations between images. V1 & PM reveal a strong correlational structure of within-epoch population responses, with comparable correlations for between-epoch blocks. Representational similarity from superficial RSP reveals a strong correlation of within-epoch population responses, with little to no correlation between-epoch blocks. Deep RSP shows little correlation between trials. All example confusion matrices, generalization matrices, and representational similarity matrices are of the same session per area.

### 2.5 Representational drift and within-sequence generalization

What is the origin of the strong encoding of information about stimulus blocks? We noticed that randomized control pre and post blocks were discriminable from each other, yet, they consisted in identical experimental protocols (i.e., images presented in the same randomized order), their only difference being the time within the session where they occurred (beginning and end of each session, respectively). This suggested that the information about stimulus blocks was not related to a difference in image transition history, but rather to a representation of the elapsed time within the session. We investigated this effect on a faster timescales by testing whether information about the passing of time was present within the sequence block, by constructing a cross-validated classifiers to decode which epoch within the sequence block an image was presented in (Fig. 4a). In all three areas (except for deep RSP), we observed significant decoding of the epoch in which an image was presented, which we interpreted as evidence of a consistent *activity drift* within a session on the timescales of minutes (Fig. 4a). We thus hypothesized that the encoding of stimulus block information may be partially due to *activity drift* occurring over the course of the recording session [Aitken et al., 2021]. While previous examples of drift showed changes over the course of days and weeks [Aitken et al., 2021, Schoonover et al., 2021a, Marks and Goard, 2021], here we investigated whether drift occurred rapidly within a single 33-minute session [Deitch et al., 2021].

We thus found the consistent presence of *activity drift* in V1, PM, and superficial RSP, defined as a significant discriminability of the epoch within a session where the same stimuli were presented. We then investigated the relationship between this activity drift and the representational geometry of image identity. We hypothesized the following two alternative scenarios for this interaction. We denoted *representational drift* the case where a drift in activity maintained the representational geometry of image identity(Fig. 4c), and a simple *activity drift* the case where changes occurred that disrupted the representational geometry.

We found evidence for *representational drift* in V1 and PM; while superficial RSP exhibited a simple *activity drift* which did not preserve representations (Fig. 4d). Let us examine more in details these findings (Fig. 4).

We investigated the evolution of the representational geometry of expected natural images under drift. Two alternative scenarios may arise: In the first scenario, the drift may occur in directions orthogonal to the stimulus-encoding ones, leading to a stable representation of image identity which can generalize across different epochs and be consistently read out by a downstream neuron, represented by a linear classifier. In the second scenario, the drift direction may overlap with the stimulus-coding axis, thus leading to epoch-specific representations which cannot generalize across epochs. We compared these two scenarios in PCA space, where we found that the representational geometry of the main sequence, defined as the relative position of responses to the ABCD images, is preserved across epochs in V1 (Fig. S6 c) and PM, but not in RSP (Fig. S6 f). This result is also replicated in the representational similarity plots of V1 and PM vs RSP, where large correlations on the off-diagonal blocks in V1 and PM suggest generalization across epochs, while the weak correlation in off-diagonal blocks in RSP suggests lack of generalization across epochs (Fig. 4e).

To quantify the difference between the two scenarios we trained linear classifiers on trials from one epoch and test their generalization performance on trials from a different epoch Bernardi et al. [2020]. We found that linearly separable image identity representations in V1 and PM, but not in RSP, generalized across epochs (Fig. 4d, Fig. S5e-h). These results show that visual representations in V1 and PM, but not RSP, maintain a consistent linear readout available to downstream neurons, providing a stable representational drift occurring on the timescale of minutes, consistent with recent studies [Aitken et al., 2021]. Visual responses in RSP, although they significantly encoded information about main sequence image identity, did not maintain the representational geometry and thus were subject to simple activity drift. Critically, these results did not depend on whether the animal was running or at rest. We performed the same decoding analyses but conditioning on trials where the animal was at rest and found the same stable representational drift in V1 and PM, but activity drift in RSP (Fig. S6).

## 3 Discussion

We found that encoding of natural scenes varies along the cortical hierarchy as well as its temporal context, namely, on the history of image-specific transitions preceding each scene (Fig. 2). Image encoding was strongly modulated by expectations, consistent with theories of hierarchical predictive coding, with increased responses to unexpected compared to expected scenes in V1 and PM at the bottom of the visual hierarchy (feedforward error propagation); while in RSP, at the top of the hierarchy, responses to unexpected scenes recapitulated the missing image (feedback predictions). Finally, we found evidence for representational drift in V1 and PM, where evoked responses maintained a faithful representational geometry despite changes in activity; but not in RSP, where activity drifts disrupted the geometry of responses. Taken together, these results suggest that visual responses in the cortical hierarchy are affected by the interaction between temporal context, representational drift, and expectations, in a way consistent with predictive coding.

We found that the stimulus block in which an image was presented in (randomized control pre and post vs early, middle, and late phases of sequence vs transition control) was encoded strongly in the neural populations of V1, PM, and even RSP (Fig. 1e, Fig. 4), the latter having similar context-dependent decoding accuracy as the other areas, despite much lower decoding performance of identities of expected images and chance-level decoding performance to unexpected images. One interpretation of this finding is that of fast representational drift occurring on the timescales of minutes, consistent with recent studies [Deitch et al., 2021]. The difference in visual representations across stimulus blocks may include other effects beyond representational drift. In particular, different blocks differ in how many images a sequence comprised (1 image in the randomized control, 2 images in the transition control, and 4 in the main sequence). History effects extending beyond pairwise transitions may thus explain the discriminability of sequence vs transition control. This was observed in sequences of randomized gabor patches [Kim et al., 2019a], but not for extended sequences of natural images. A recent study suggested that representational drift might be explained away by changes in an animal’s behavioral state (e.g., running vs. resting) occurring within a single recording session Sadeh and Clopath [2022]. However, we found that representational drift persisted even after controlling for changes in an animal’s behavioral state (Fig. S6), suggesting that it may represent an important feature of cortical dynamics.

The transition between successive flashed images is an essential part of the encoding of naturalistic stimuli in 2-photon datasets. Past studies from the Allen Institute have reported responses to naturalistic images that are presented in random order [de Vries et al., 2020, Siegle et al., 2021]. Although the decoding of image identity was significantly stronger in electrophysiological recordings, we showed that our results are consistent with calcium imaging recordings from those publicly available datasets (Fig. S1). Other studies have reported responses to naturalistic images that are presented with an interleaving gray screen [Kowalewski et al., 2021], thus neglecting information about transitions between scenes encoding stimulus responses dynamics. Previous studies suggested that only the preceding image matters for distinguishing between sequential image presentations [Nikolić et al., 2009]. However, in V1 and PM, we found a significant increase in decoding performance of unexpected images in the four-image sequence compared to randomized control and two-image transition control blocks, highlighting the potential role of longer sequences in enhancing encoding of visual stimuli.

Our results on representational drift are consistent with previous studies in V1 and PM [Deitch et al., 2021, Marks and Goard, 2021, Aitken et al., 2021], and extend them to RSP. Our measure of representational drift time decoding and cross-epoch generalization performance allows us to simultaneously observe whether drift is occurring, but it also allows us to test for generalization in the population response. This idea is fundamentally linked to the problem of representational drift: Can neural populations faithfully encode for persistent representations while being subject to drift in activity? We found that representations of expected natural images in V1 and PM generalize despite drift, while RSP does not, suggesting that representational drift features may be area-specific.

We found that visual responses along the hierarchy were strongly affected by expectations in a way that is consistent with the classic theory of hierarchical predictive coding [Rao and Ballard, 1999]. We investigated this effect by examining differences in decoding performances of expected vs unexpected natural scenes along the visual hierarchy. First, we found that population responses in V1 and PM, but not RSP, significantly encoded unexpected (oddball) images identity across all stimulus blocks (Fig. 2h). Moreover, single cell selectivity for oddball images was stronger than for the main sequence images ABCD in V1 and PM (Fig. 1d). Furthermore, unexpected stimuli were encoded significantly better in the sequence block than in both the transition ctrl and the randomized ctrl for both V1 and PM (Fig. 2h). We interpret this finding within the paradigm of predictive coding, following recent experimental evidence in [Gillon et al., 2021]. In the sequence block, violations of the expected image D cause prediction error signals that are not present in the transition control block, which shuffles the expected sequence of images into a random set of pairwise images. No expectation can be formed in the transition control beyond the pairwise transition.

Combining expected and unexpected stimuli paradigms into one classification task (Fig. 3), we found that V1 and PM significantly encoded information about both expected (main-sequence) vs unexpected (oddball) images. In RSP, population responses to the unexpected oddball images were indistinguishable from responses to the expected, but missing, image D (Fig. 3c). One interpretation of this finding is that RSP encodes for the expectation of the missing image, rather than what was actually shown. Indeed, we found that RSP responses do not encode for distinct oddball representations at all (Fig. 2g) but rather confound oddball responses with the missing D, but not A,B, or C. Our results suggest that expectation may play a role in RSP, consistent with hierarchical predictive coding theory which posits higher cortical areas to encode prediction [Rao and Ballard, 1999, Keller et al., 2012].

## 4 Methods

### 4.1 Experimental data

All experiments and procedures were performed in accordance with protocols approved by the Allen Institute Animal Care and Use Committee. The dataset used in this paper was collected as part of the Allen Institute for Brain Science’s OpenScope initiative. Data were collected and processed using the Allen Brain Observatory data collection and processing pipelines [de Vries et al., 2020]. Here we include a brief description on experimental procedure. The full details on the data collection and process are described in [de Vries et al., 2020].

Transgenic mice expressing GCaMP6f in excitatory cells were used (Slc17a7-IRES2-Cre x CaMk2-tTA x Ai93(GCaMP6f). Only sessions that pass quality control criteria described in [de Vries et al., 2020] were included in our analyses, resulting in total N = 14 mice, with 16 sessions from V1 (2299 neurons), 23 sessions from PM (2071 neurons), and 12 sessions from RSP (1628 neurons). Two-photon calcium imaging was performed a Nikon A1R MP+, with imaging depths ranging 125 - 450*µm* to capture neuronal activities across cortical layers.

Mice were injected with a retrograde tracer (AAVRetro.CAG.mRuby3) in either V1 or RSP, however this data was not used in our study. No differences were observed in labeled vs unlabeled cells for any of the quantifications in this manuscript. Information on the identity of retrogradely labeled cells is available upon request.

Each mouse experienced 4 imaging sessions, each with the same stimulus protocol. Two sessions were recorded in one cortical area (ex: V1, PM, or RSP) at 2 cortical depths, typically around 175*µm* (approximately layer 2/3) and 375*µm* (approximately layer 5), and two sessions were recorded at similar depths in a different cortical area. Depth was chosen based on cre-line analysis performed in [de Vries et al., 2020] and takes into account brain compression due to implant and off-normal imaging axis. Areas were chosen such that the injection site for retrograde labeling was never imaged. For the full details on on animal surgery, habituation, quality control, data collection, and post-collection data processing, please see [de Vries et al., 2020].

The dataset along with the code for analyses included in this paper is available at https://github.com/AllenInstitute/openscope_temporal_context, and the full dataset is publicly available in Neurodata Without Borders (NWB) format in the DANDI Archive Lecoq et al. [2023].

### 4.2 Stimulus protocol

Visual stimuli consisted of a subset of the natural images publicly available Allen Brain Observatory dataset (https://observatory.brain-map.org/visualcoding/;[de Vries et al., 2020]). The images were presented in grayscale, contrast normalized, matched to have equal mean luminance, and resized to 1,174 × 918 pixels. Four natural images (Brain Observatory image IDs: im013, im026, im068, im078) were used to form a familiar sequence of four images (ABCD), and ten additional images served as unexpected oddball images (im06, im017, im022, im051, im071, im089, im103, im110, im111, im112).

Stimuli were presented in 4 distinct blocks over the course of a 1 hour imaging session (64 minutes). Individual stimuli were presented for 250ms with no intervening gray period (i.e. one image after the other) in all blocks. Blocks were separated by 60 seconds of gray screen in which spontaneous activity could be measured. A schematic of the stimulus design is shown in Fig. 1b.

In the Randomized Control blocks, the 14 images (4 sequence images and 10 oddball images) were presented in random order. Each image was shown for 250ms with no intervening gray screen. Each image was presented 30 times. The randomized control stimulus block was presented once at the beginning of the session and once at the end of the session and lasted 0.25s x 30 repeats = 105 seconds each time.

In the Sequence block, the series of expected sequence images ABCD was repeated 20 times per cycle, with an oddball image randomly taking the place of image D after 10-19 repeats of the sequence in that cycle. The number of sequence repeats between oddball image occurrences was random and not predictable. Each oddball image was presented a total of 10 times throughout the sequence block, for a total of 100 cycles (10 images x 10 cycles each). Each main sequence image ABCD was presented 20 times per cycle x 100 cycles = 2000 times. The entire block lasted 33.33 minutes.

In the Transition Control block, the image transitions shown in the Sequence block were preserved as pairs (AB, BC, CD, DA, CX*_i_*, X*_i_*A, with *i* = 1*, …,* 10 oddball images, giving a total of 24 pairs), but the global sequence was not preserved. Each image transition pair was treated as a distinct stimulus (lasting 0.5 seconds) and presented in random order. Each image pair was shown 30 times, for a total of 24 pairs x 30 repeats = 720 pair presentations. The Transition Control block was 6 minutes in duration (720 pair presentations x 0.5 seconds per pair = 360 seconds).

Two additional stimuli not used in this study were also shown. The occlusion stimulus, consisted of the 10 oddball images with 6 differing levels of spatial occlusion. Each occlusion image was presented for 0.5 seconds, with 0.5 seconds of gray screen between stimuli. Each occlusion image was presented 10 times, for a total of 10 images x 6 occlusion levels x 10 repeats = 600 individual stimulus presentations, for a total of 600 seconds = 6 minutes. A 30 second natural movie clip (Brain Observatory stimulus set Natural Movie 1) was repeated 10 times at the end of the session.

### 4.3 Decoding analysis

We decoded stimulus block, image identity, and time within a session from single-trial population response vectors. To construct these response vectors, we performed deconvolution [Jewell et al., 2019, de Vries et al., 2020] on the delta fluorescence traces and took the deconvolved events within a circumscribed response window of 50ms to 250ms relative to stimulus onset. These response vectors were first flattened and passed through principle components analysis before being used in the decoding analysis. Specifically, we trained multi-class linear support vector machine (SVM) classifiers to decode the above categorical variables from the single-trial population response vectors. Stratified 5-fold cross-validation was used for decoding expected images (Fig. 2d), block and “time” (Fig. 4), ensuring equal number of trials for each class. When performing classification with oddball images (Fig. 2h and Fig. 3), 10-fold stratified cross-validation was performed, as each oddball was only presented 10 times during the sequence stimulus block.

To calculate the statistical significance of decoding accuracies, we performed an iterative shuffle procedure on each fold of the cross-validation. In each shuffle, the training labels which the classifer was trained to decode were shuffled randomly across trials of the training set, and the classifer’s accuracy was evaluated on the unshuffled test data-set. This shuffle was performed 100 times to create a shuffle distribution of decoding accuracies for each fold of the cross-validation. From these distributions we calculated the z-score of decoding accuracy for each class in each cross-validation fold. These z-scores were then averaged across the folds of cross-validation and used to calculate the overall p-value of the decoding accuracy obtained on the original data.

### 4.4 Validation

To examine issues of stimulus design or recording methodology, we performed the same decoding analysis on three independent datasets where natural images were presented in random order. First we compared decoding performances evaluated on data from the publicly available Allen Brain Observatory Visual Coding (“ophys”) and Neuropixels (“ephys”) datasets [de Vries et al., 2020, Siegle et al., 2019]. Both experiments consisted of 118 natural image stimuli from the same dataset as ours with identical presentation protocol. We confirmed significant decoding accuracy across all 118 images using electrophysiological population responses reliably across experiments (Fig. S1g). Using population responses extracted from 2-photon imaging data also showed significant decoding performance on image identification, but at a much reduced level (Fig. S1h). Restricting ourselves to the 14 images presented in our experiment (4 main-sequence and 10 oddball images) increased decoding performance for 2-photon data.

### 4.5 Single-cell selectivity

To assess single cell selectivity to particular natural images and stimulus block, we performed simple one-way anovas using the open source python software pingouin [Vallat, 2018]. For each stimulus block, we determined if a cell was selective to one of the four main-sequence images or to one of the 10 oddball images by using the population response vectors for those trials in which the images were presented (Fig. 1d). We ensured each group had an equal number of trials. For each image, we determined if a cell was selective to the block in which it was presented in by using population response vectors for the trials in which the same image was presented in different blocks (Fig. 1e). P-values were calculated from the F-distribution and were considered significant (selective) if below a threshold of 0.05.

## Acknowledgments

The data presented herein were obtained at the Allen Brain Observatory as part of the OpenScope project, which is operated by the Allen Institute for Brain Science. This work was supported by the Allen Institute and in part by the Falconwood Foundation. We thank Allan Jones for providing the critical environment that enabled our large-scale team effort. We thank the Allen Institute founder, Paul G Allen, for his vision, encouragement, and support. Research reported in this publication was supported by the National Eye Institute of the National Institutes of Health under Award Number K99/R00 EY030840 to H.C. DGW and LM are partially supported by NINDS: R01-NS118461 (BRAIN Initiative), NIDA: R01-DA055439 (NSF CRCNS). The content is solely the responsibility of the authors and does not necessarily represent the official views of the National Institutes of Health.

## Author contributions

See Supplementary Figure S7.

## Declaration of interests

The authors declare no competing interests.

---

**Figure S1.**
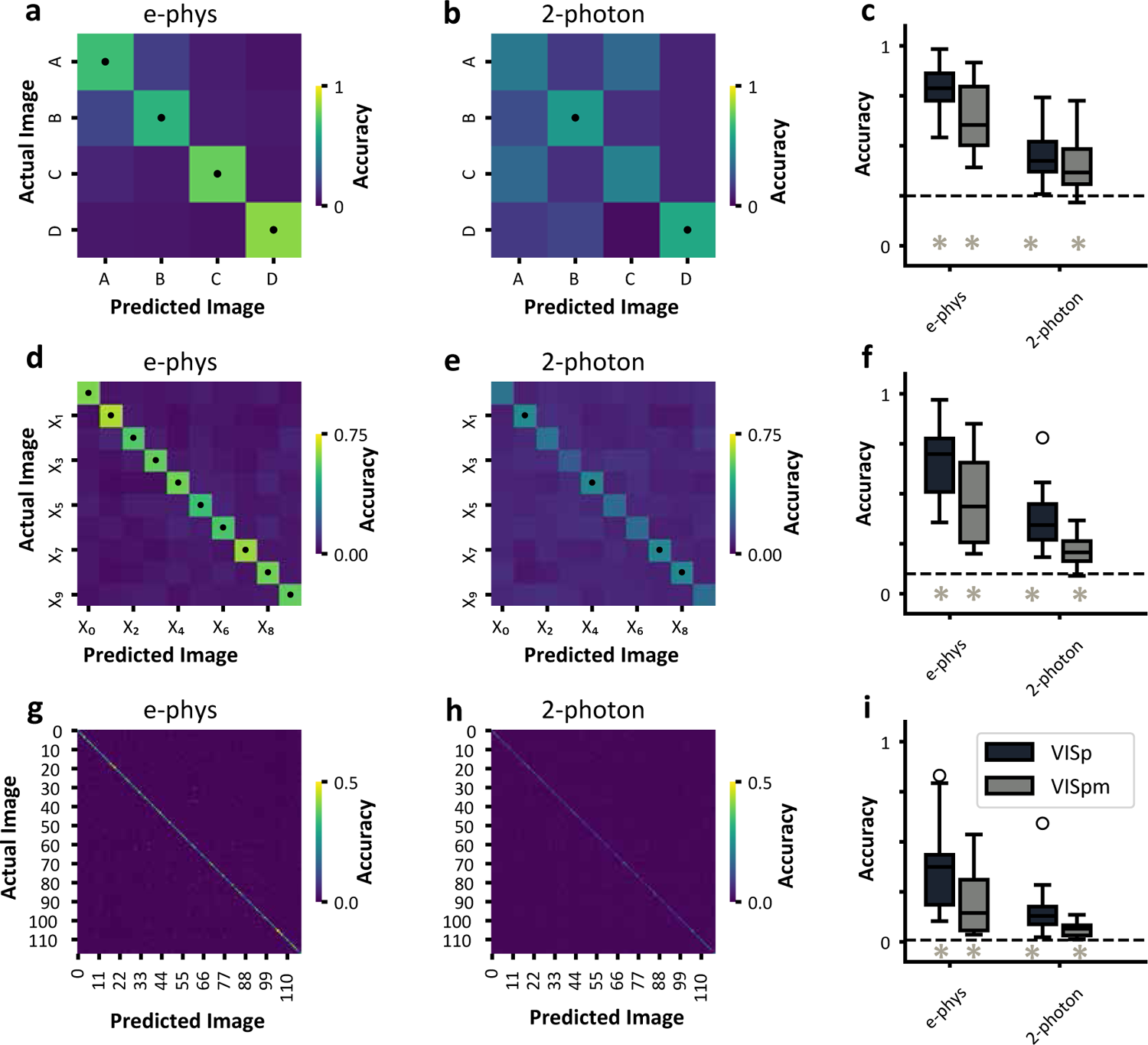
Validation of decoding results with electrophysiological [Siegle et al., 2021] and two-photon [de Vries et al., 2020] functional datasets from mouse V1 and PM a) Example confusion matrix from one session of V1 e-phys data trained to classify the four main-sequence images within our study. b) Example confusion matrix from one session of V1 two-photon data trained to classify the four main-sequence images within our study. c) Summary boxplot over 32 sessions of e-phys and 39 sessions of 2-photon data. d - f) Same as a-c, but for the 10 oddball images. g - i) Same as a-c, but for all 118 images within the Berkeley Segmentation Dataset

**Figure S2.**
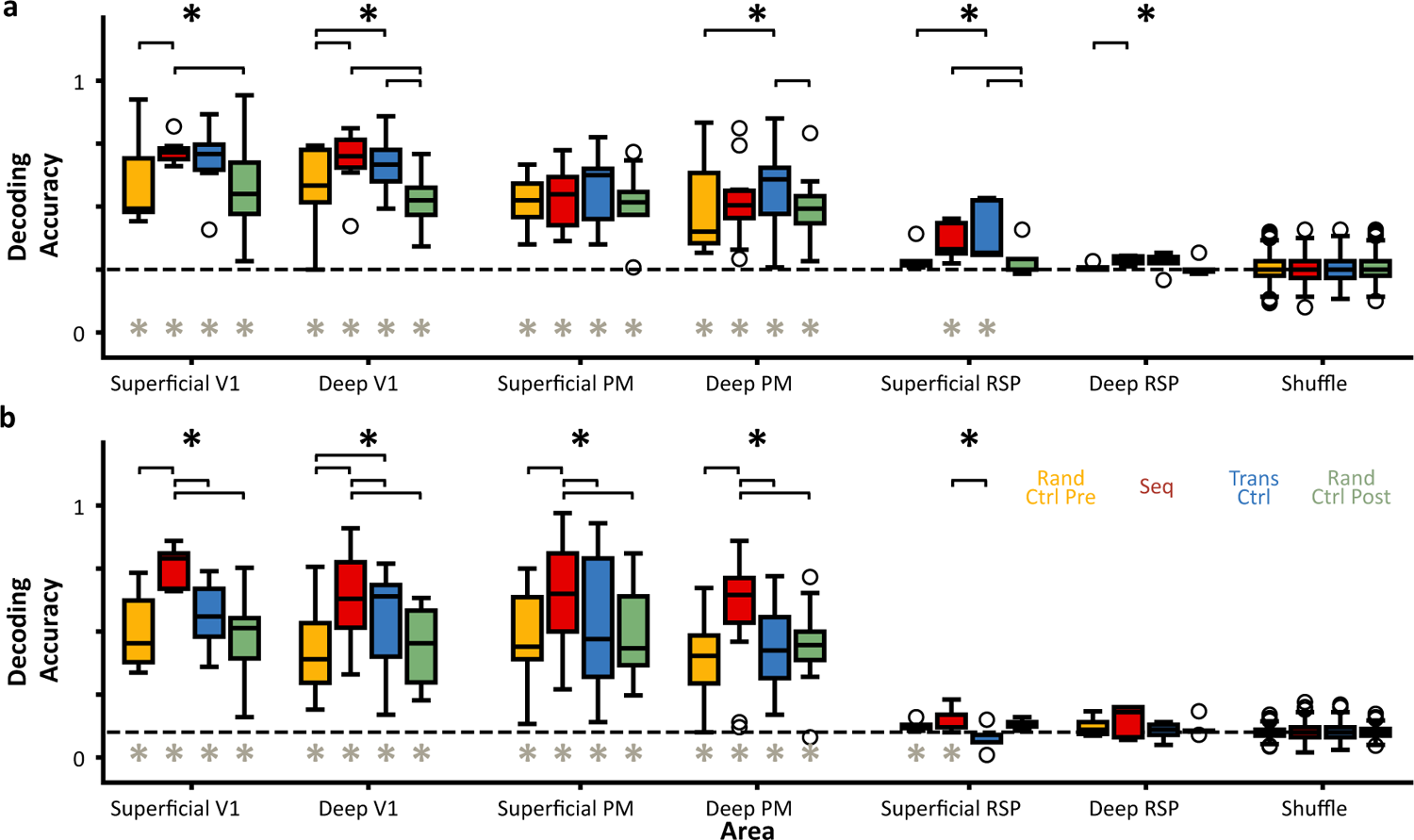
Decoding of natural images across different stimulus blocks, separated by area and depth. **a & b)** Summary boxplots show the decoding performance of main sequence images (in *a*) and unexpected oddball images (in *b*) across all stimulus blocks for each unique area and depth.

**Figure S3.**
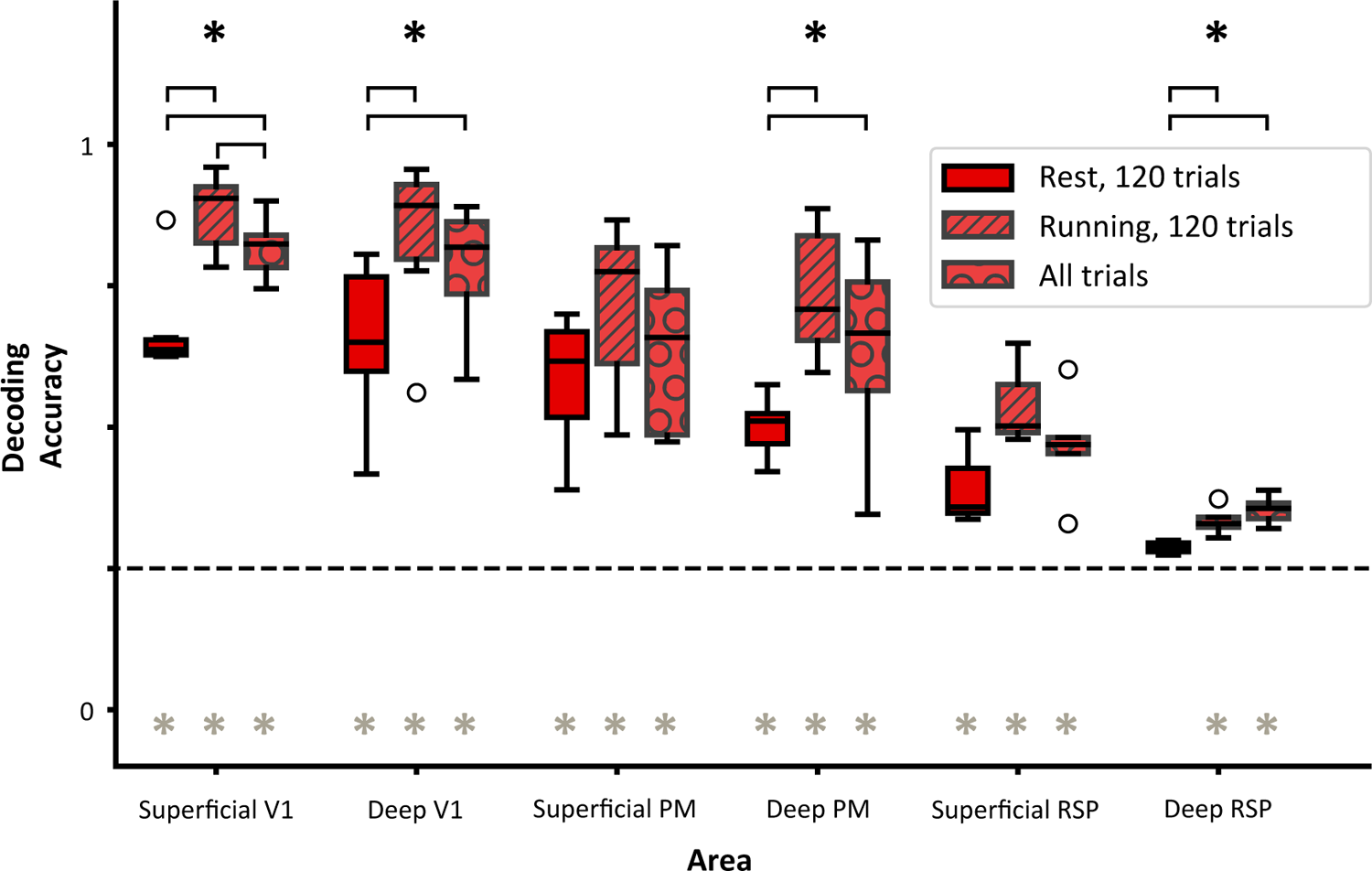
Decoding of expected natural images is modulated by running. In the sequence stimulus block, classifiers were constructed using either the first 120 trials (30 trials per image) where the animal was at rest, or the first 120 trials where the animal was running, or using all trials (1900 trials per image) in the sequence block. The decoding accuracy of such classifiers varied significantly in all areas.

**Figure S4.**
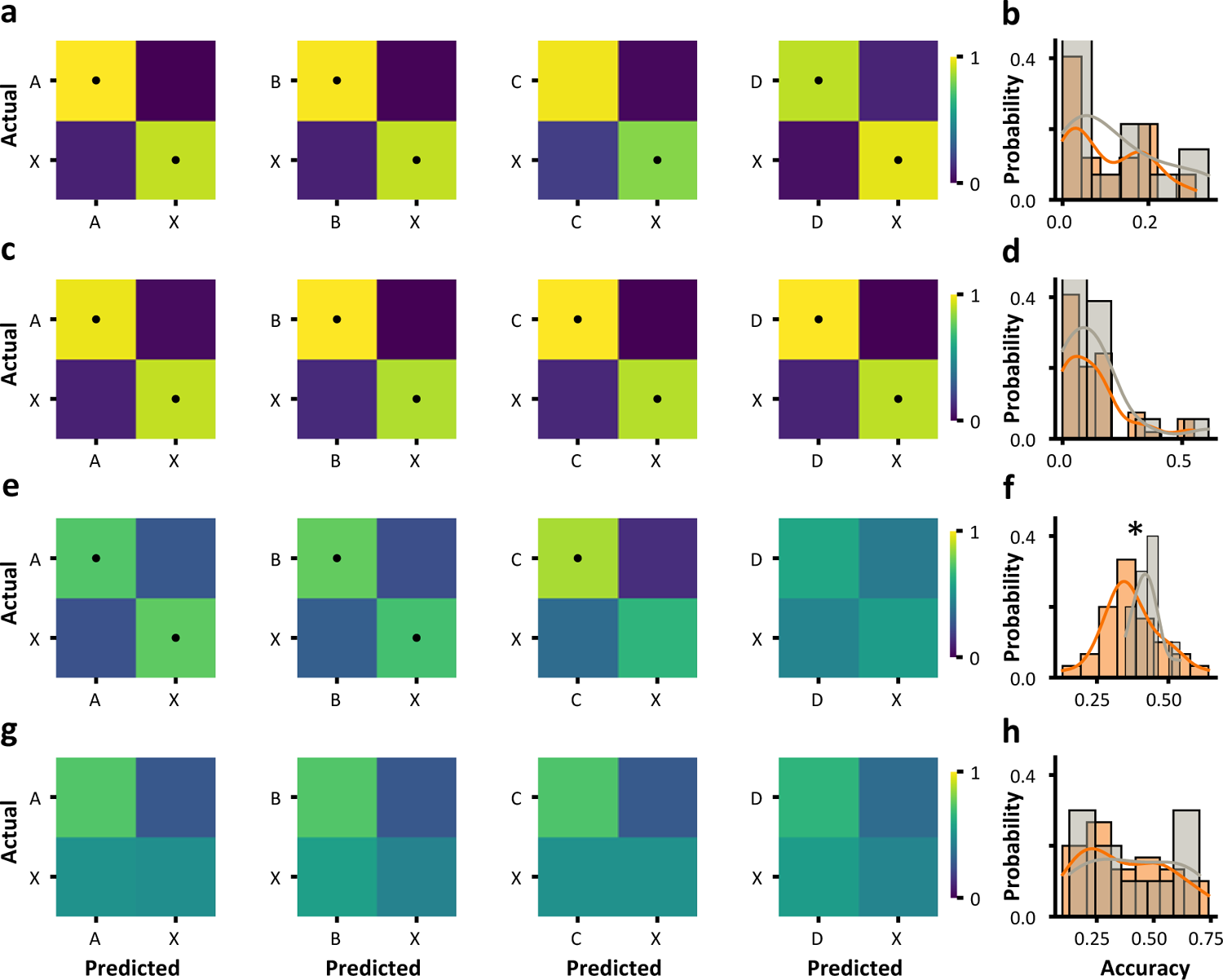
Decoding responses to expected and unexpected natural images reveals possible predictive coding mechanism in RSP. **a)** Four separate classifiers constructed to decode each expected main sequence image from the unexpected oddball images. This first row is for superficial V1, which recapitulates the main Fig. 3e results. **b)** Histogram of the relative miss rates of main sequence images (A instead of X, B instead of X, C instead of X) compared to that misclassifying images D and X. Superficial V1 shows no significant difference in the DX miss-rate relative to the ABCD miss-rate. **c-d)** Same as first row, but for superficial PM. **e-f)** Same as first row, but for superficial RSP. **f)** Superficial RSP reveals a significant increase in the DX miss-rate relative to the ABCD miss-rate. **g-h)** Same as first row, but for deep RSP.

**Figure S5.**
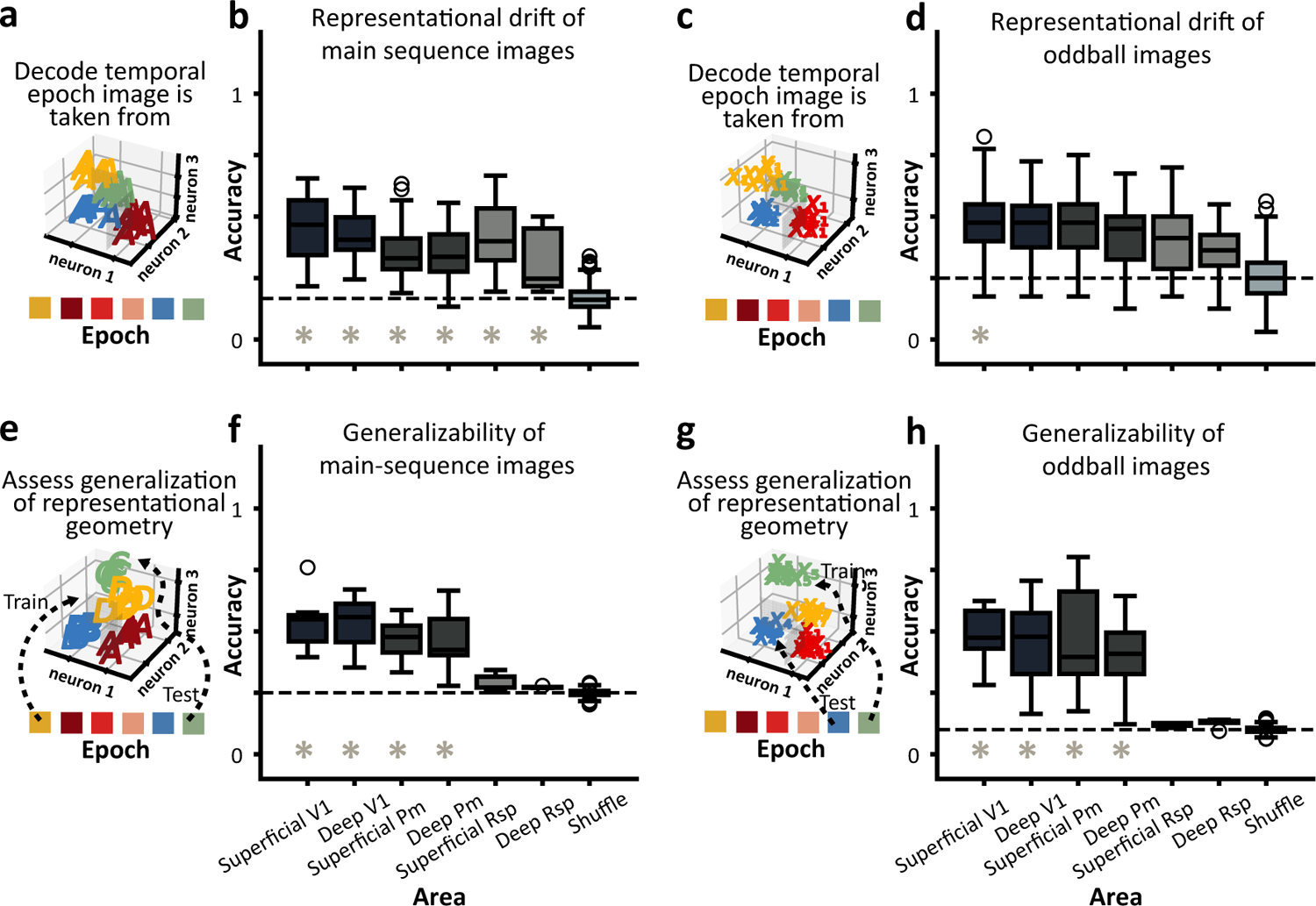
Generalization under representational drift, separated by area and depth. **a)** Schematic showing linearly separable population responses to the same main sequence natural image presented in different stimulus blocks. **b)** Summary boxplots show significant representational drift of main sequence images across areas. **c - d)** Same as a & b, but for unexpected oddball images. **e)** Schematic showing population responses to main sequence images in different epochs of the session, with training and testing occurring using data from different stimulus blocks (epochs). **f)** Summary boxplots show significant generalizability of main sequence images in V1 and PM, but not in RSP. **g - h)** Same as e & f, but for unexpected oddball images.

**Figure S6.**
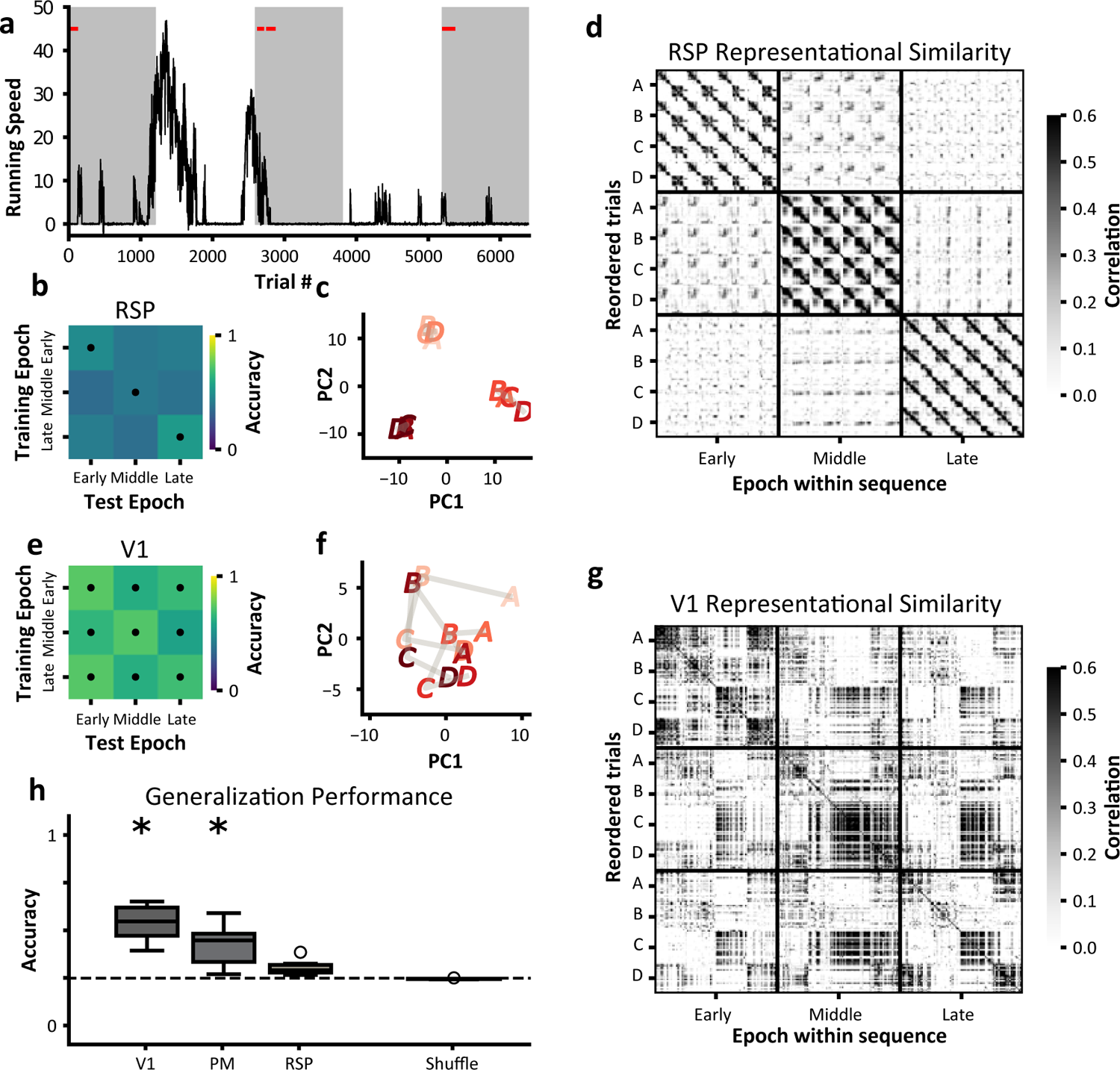
Generalization performance under representational drift, whilst controlling for behavioral state. **a)** Example running speed trace of an animal, where the early, middle, and late epochs are defined by gray highlights. The first 30 trials per image where the animal was at rest were taken to perform the decoding analysis. Rest is defined as less than 1 cm/s. **b)** Confusion matrix from RSP shows that only within-epoch decoding is possible (main diagonal). **c)** PCA of main sequence PSTHs from RSP show epochs cluster apart. **d)** Representational similarity from RSP reveals a strong correlation of within-epoch population responses, with little to no correlation between-epoch blocks. **e)** Confusion matrix from V1 shows that both within-epoch decoding (main diagonal) and between-epoch decoding (off-diagonal) is possible. **f)** PCA of main sequence PSTHs from V1 shows conservation of geometry between epochs. **g)** Representational similarity from V1 reveals a strong correlation of within-epoch population responses, with comparable correlations for between-epoch blocks. **h)** Generalization decoding results over single sessions.

**Figure S7.**
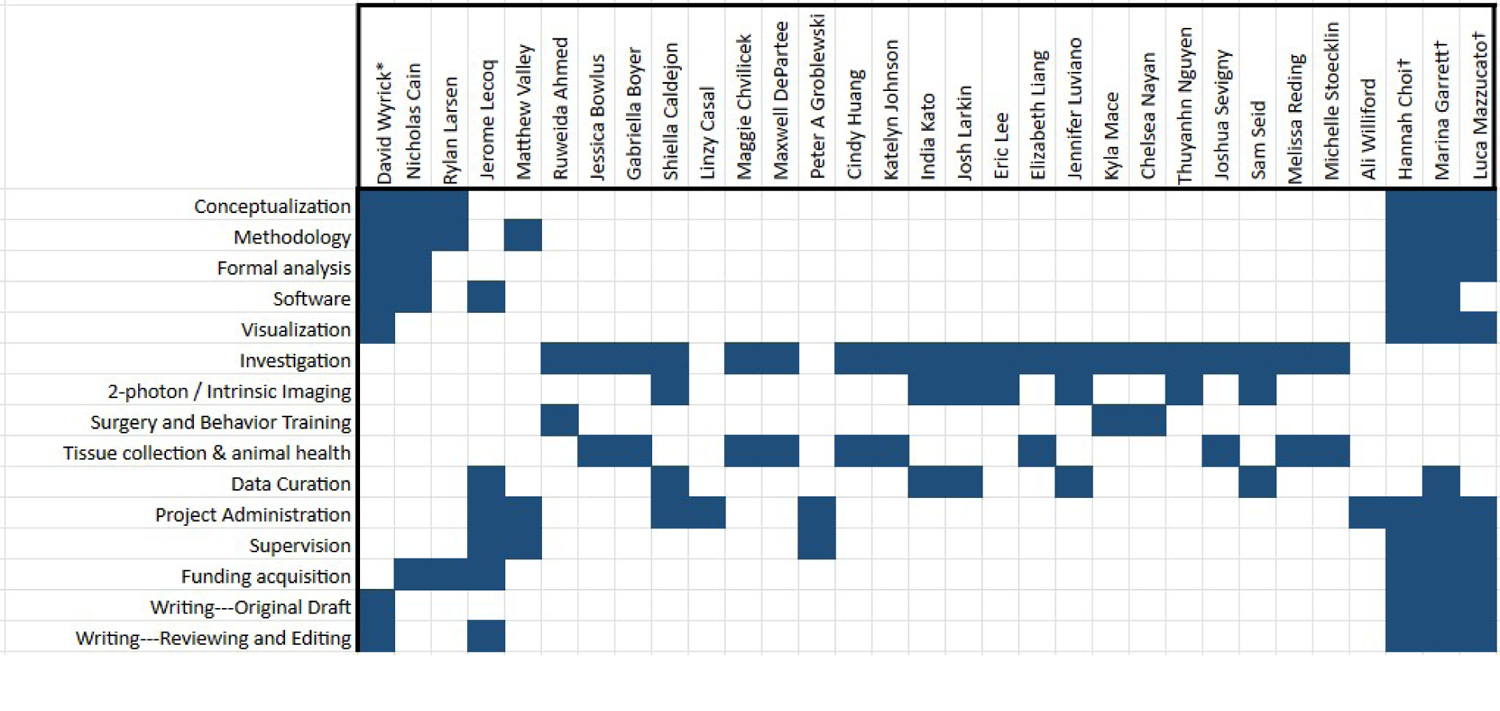
Author Contributions

## Notes

https://dandiarchive.org/dandiset/000488/draft

### Competing Interest Statement

The authors have declared no competing interest.

### Summary of Updates

Author list correction

## References

K. Aitken, M. Garrett, S. Olsen, and S. Mihalas. The Geometry of Representational Drift in Natural and Artificial Neural Networks. bioRxiv, page 2021.12.13.472494, 2021. URL https://www.biorxiv.org/content/10.1101/2021.12.13.472494v2%, https://www.biorxiv.org/content/10.1101/2021.12.13.472494v2.abstract.

K. Aitken, M. Garrett, S. Olsen, and S. Mihalas. The geometry of representational drift in natural and artificial neural networks. PLOS Computational Biology, 18(11):e1010716, Nov. 2022. ISSN 1553-7358. doi: 10.1371/journal.pcbi.1010716. URL https://dx.plos.org/10.1371/journal.pcbi.1010716.

A. S. Alexander and D. A. Nitz. Retrosplenial cortex maps the conjunction of internal and external spaces. Nature Neuroscience, 18(8):1143–1151, 2015. ISSN 15461726. doi: 10.1038/nn.4058.

A. S. Alexander, R. Place, M. J. Starrett, E. R. Chrastil, and D. A. Nitz. Rethinking retrosplenial cortex: Perspectives and predictions. Neuron, page S0896627322010273, Dec. 2022. ISSN 08966273. doi: 10.1016/j.neuron.2022.11.006. URL https://linkinghub.elsevier.com/retrieve/pii/S0896627322010273.

A. Bastos, W. Usrey, R. Adams, G. Mangun, P. Fries, and K. Friston. Canonical microcircuits for predictive coding. Neuron, 76(4):695–711, 2012. ISSN 0896-6273. doi: https://doi.org/10.1016/j.neuron.2012.10.038. URL https://www.sciencedirect.com/science/article/pii/S0896627312009592.

S. Bernardi, M. K. Benna, M. Rigotti, J. Munuera, S. Fusi, and C. D. Salzman. The Geometry of Abstraction in the Hippocampus and Prefrontal Cortex. Cell, 183(4):954–967.e21, 2020. ISSN 10974172. doi: 10.1016/j.cell.2020.09.031.

S. E. de Vries, J. A. Lecoq, M. A. Buice, P. A. Groblewski, G. K. Ocker, M. Oliver, D. Feng, N. Cain, P. Ledochowitsch, D. Millman, K. Roll, M. Garrett, T. Keenan, L. Kuan, S. Mihalas, S. Olsen, C. Thompson, W. Wakeman, J. Waters, D. Williams, C. Barber, N. Berbesque, B. Blanchard, N. Bowles, S. D. Caldejon, L. Casal, A. Cho, S. Cross, C. Dang, T. Dolbeare, M. Edwards, J. Galbraith, N. Gaudreault, T. L. Gilbert, F. Griffin, P. Hargrave, R. Howard, L. Huang, S. Jewell, N. Keller, U. Knoblich, J. D. Larkin, R. Larsen, C. Lau, E. Lee, F. Lee, A. Leon, L. Li, F. Long, J. Luviano, K. Mace, T. Nguyen, J. Perkins, M. Robertson, S. Seid, E. Shea-Brown, J. Shi, N. Sjoquist, C. Slaughterbeck, D. Sullivan, R. Valenza, C. White, A. Williford, D. M. Witten, J. Zhuang, H. Zeng, C. Farrell, L. Ng, A. Bernard, J. W. Phillips, R. C. Reid, and C. Koch. A large-scale standardized physiological survey reveals functional organization of the mouse visual cortex. Nature Neuroscience, 23(1):138–151, 2020. ISSN 15461726. doi: 10.1038/s41593-019-0550-9.

D. Deitch, A. Rubin, and Y. Ziv. Representational drift in the mouse visual cortex. Current Biology, 31(19):4327–4339.e6, 2021. ISSN 18790445. doi: 10.1016/j.cub.2021.07.062. URL https://doi.org/10.1016/j.cub.2021.07.062.

L. N. Driscoll, N. L. Pettit, M. Minderer, S. N. Chettih, and C. D. Harvey. Dynamic Reorganization of Neuronal Activity Patterns in Parietal Cortex. Cell, 170(5):986–999.e16, Aug. 2017. ISSN 10974172. doi: 10.1016/j.cell.2017.07.021. URL https://linkinghub.elsevier.com/retrieve/pii/S0092867417308280. Publisher: Elsevier Inc. ISBN: 1097-4172 (Electronic) 0092-8674 (Linking).

L. N. Driscoll, L. Duncker, and C. D. Harvey. Representational drift: Emerging theories for continual learning and experimental future directions. Current Opinion in Neurobiology, 76: 102609, Oct. 2022. ISSN 1873-6882. doi: 10.1016/j.conb.2022.102609.

K. Friston. A theory of cortical responses. Philosophical Transactions of the Royal Society B: Biological Sciences, 360(1456):815–836, 2005. ISSN 09628436. doi: 10.1098/rstb.2005.1622.

J. P. Gavornik and M. F. Bear. Learned spatiotemporal sequence recognition and prediction in primary visual cortex. Nature neuroscience, 17(5):732–737, 2014.

C. J. Gillon, J. E. Pina, J. A. Lecoq, R. Ahmed, Y. N. Billeh, S. Caldejon, P. Groblewski, T. M. Henley, E. Lee, J. Luviano, et al. Learning from unexpected events in the neocortical microcircuit. BioRxiv, 2021.

J. A. Harris, S. Mihalas, K. E. Hirokawa, J. D. Whitesell, H. Choi, A. Bernard, P. Bohn, S. Caldejon, L. Casal, A. Cho, A. Feiner, D. Feng, N. Gaudreault, C. R. Gerfen, N. Graddis, P. A. Groblewski, A. M. Henry, A. Ho, R. Howard, J. E. Knox, L. Kuan, X. Kuang, J. Lecoq, P. Lesnar, Y. Li, J. Luviano, S. McConoughey, M. T. Mortrud, M. Naeemi, L. Ng, S. W. Oh, B. Ouellette, E. Shen, S. A. Sorensen, W. Wakeman, Q. Wang, Y. Wang, A. Williford, J. W. Phillips, A. R. Jones, C. Koch, and H. Zeng. Hierarchical organization of cortical and thalamic connectivity. Nature, 575(7781):195–202, 2019. ISSN 14764687. doi: 10.1038/s41586-019-1716-z. URL http://dx.doi.org/10.1038/s41586-019-1716-z. Publisher: Springer US.

J. Homann, S. A. Koay, K. S. Chen, D. W. Tank, and M. J. Berry. Novel stimuli evoke excess activity in the mouse primary visual cortex. Proceedings of the National Academy of Sciences, 119(5):e2108882119, 2022.

D. H. Hubel and T. N. Wiesel. Receptive fields of single neurones in the cat’s striate cortex. The Journal of physiology, 148(3):574, 1959.

D. H. Hubel and T. N. Wiesel. Receptive fields, binocular interaction and functional architecture in the cat’s visual cortex. The Journal of physiology, 160(1):106, 1962.

M. Ito and C. D. Gilbert. Attention modulates contextual influences in the primary visual cortex of alert monkeys. Neuron, 22(3):593–604, 1999.

S. W. Jewell, T. D. Hocking, P. Fearnhead, and D. M. Witten. Fast nonconvex deconvolution of calcium imaging data. Biostatistics, Feb. 2019. doi: 10.1093/biostatistics/kxy083. URL https://doi.org/10.1093/biostatistics/kxy083.

A. Jezzini, L. Mazzucato, G. La Camera, and A. Fontanini. Processing of hedonic and chemosensory features of taste in medial prefrontal and insular networks. Journal of Neuroscience, 33(48):18966–18978, 2013.

G. B. Keller and T. D. Mrsic-Flogel. Predictive processing: A canonical cortical computation. Neuron, 100(2):424–435, 2018. ISSN 0896-6273. doi: https://doi.org/10.1016/j.neuron.2018.10.003. URL https://www.sciencedirect.com/science/article/pii/S0896627318308572.

G. B. Keller, T. Bonhoeffer, and M. Hübener. Sensorimotor mismatch signals in primary visual cortex of the behaving mouse. Neuron, 74(5):809–815, 2012.

A. G. Khan, J. Poort, A. Chadwick, A. Blot, M. Sahani, T. D. Mrsic-Flogel, and S. B. Hofer. Distinct learning-induced changes in stimulus selectivity and interactions of gabaergic interneuron classes in visual cortex. Nature neuroscience, 21(6):851–859, 2018.

H. Kim, J. Homann, D. Tank, and M. Berry. A Long Timescale Stimulus History Effect in the Primary Visual Cortex. bioRxiv, 2019a. doi: 10.1101/585539.

H. Kim, J. Homann, D. W. Tank, and M. J. Berry. A long timescale stimulus history effect in the primary visual cortex. BioRxiv, page 585539, 2019b.

N. N. Kowalewski, J. Kauttonen, P. L. Stan, B. B. Jeon, T. Fuchs, S. M. Chase, T. S. Lee, and S. J. Kuhlman. Development of Natural Scene Representation in Primary Visual Cortex Requires Early Postnatal Experience. Current Biology, 31(2):369–380.e5, 2021. ISSN 18790445. doi: 10.1016/j.cub.2020.10.046. URL https://doi.org/10.1016/j.cub.2020.10.046.

J. A. Lecoq, M. Garrett, H. Choi, L. Mazzucato, and D. Wyrick. Allen institute openscope - differential encoding of temporal context and expectation (version 0.230602.2022) [data set]. DANDI archive, 2023. doi: 10.48324/dandi.000488/0.230602.2022. URL https://doi.org/10.48324/dandi.000488/0.230602.2022.

H. Makino and T. Komiyama. Learning enhances the relative impact of top-down processing in the visual cortex. 18(8), 2015. doi: 10.1038/nn.4061.

T. D. Marks and M. J. Goard. Stimulus-dependent representational drift in primary visual cortex. Nature Communications, 12(1):1–16, 2021. ISSN 20411723. doi: 10.1038/s41467-021-25436-3. URL http://dx.doi.org/10.1038/s41467-021-25436-3.

C. McAdams and R. Reid. Attention modulates the responses of simple cells in monkey primary visual cortex. Journal of Neuroscience, 25(47):11023–11033, 2005. doi: 10.1523/jneurosci.2904-05.2005.

M. J. McGinley, M. Vinck, J. Reimer, R. Batista-Brito, E. Zagha, C. R. Cadwell, A. S. Tolias, J. A. Cardin, and D. A. McCormick. Waking state: rapid variations modulate neural and behavioral responses. Neuron, 87(6):1143–1161, 2015.

D. B. McMahon and C. R. Olson. Repetition suppression in monkey inferotemporal cortex: relation to behavioral priming. Journal of neurophysiology, 97(5):3532–3543, 2007.

T. Meyer, S. Ramachandran, and C. R. Olson. Statistical learning of serial visual transitions by neurons in monkey inferotemporal cortex. Journal of Neuroscience, 34(28):9332–9337, 2014.

T. Murakami, T. Yoshida, T. Matsui, and K. Ohki. Wide-field Ca2+ imaging reveals visually evoked activity in the retrosplenial area. Frontiers in Molecular Neuroscience, 08(June):1–12, 2015. ISSN 1662-5099. doi: 10.3389/fnmol.2015.00020. URL http://journal.frontiersin.org/article/10.3389/fnmol.2015.00020.

S. Musall, M. T. Kaufman, A. L. Juavinett, S. Gluf, and A. K. Churchland. Single-trial neural dynamics are dominated by richly varied movements. bioRxiv, page 308288, 2019.

D. B. Nestvogel and D. A. McCormick. Visual thalamocortical mechanisms of waking state-dependent activity and alpha oscillations. Neuron, 110(1):120–138, 2022.

C. M. Niell and M. P. Stryker. Modulation of visual responses by behavioral state in mouse visual cortex. Neuron, 65(4):472–479, 2010.

D. Nikolić, S. Häusler, W. Singer, and W. Maass. Distributed fading memory for stimulus properties in the primary visual cortex. PLoS biology, 7(12):e1000260, 2009.

J. Poort, A. G. Khan, M. Pachitariu, A. Nemri, I. Orsolic, J. Krupic, M. Bauza, M. Sahani, G. B. Keller, T. D. Mrsic-Flogel, and S. B. Hofer. Learning Enhances Sensory and Multiple Non-sensory Representations in Primary Visual Cortex. Neuron, 86(6):1478–1490, June 2015. ISSN 08966273. doi: 10.1016/j.neuron.2015.05.037. URL http://linkinghub.elsevier.com/retrieve/pii/S0896627315004766. Publisher: The Authors.

J. Poort, K. A. Wilmes, A. Blot, A. Chadwick, M. Sahani, C. Clopath, T. D. Mrsic-Flogel, S. B. Hofer, and A. G. Khan. Learning and attention increase visual response selectivity through distinct mechanisms. Neuron, page S0896627321009545, Dec. 2021. ISSN 08966273. doi: 10.1016/j.neuron.2021.11.016. URL https://linkinghub.elsevier.com/retrieve/pii/S0896627321009545.

S. Qin, S. Farashahi, D. Lipshutz, A. M. Sengupta, D. B. Chklovskii, and C. Pehlevan. Coordinated drift of receptive fields in hebbian/anti-hebbian network models during noisy representation learning. Nature Neuroscience, pages 1–11, 2023.

M. Ramadan, E. K. Lee, S. de Vries, S. Caldejon, K. Roll, F. Griffin, T. V. Nguyen, J. Larkin, P. Rhoads, K. Mace, et al. A standardized nonvisual behavioral event is broadcasted homogeneously across cortical visual areas without modulating visual responses. Eneuro, 2022.

R. P. Rao and D. H. Ballard. Predictive coding in the visual cortex: a functional interpretation of some extra-classical receptive-field effects. Nature neuroscience, 2(1):79–87, 1999.

M. E. Rule, T. O’Leary, and C. D. Harvey. Causes and consequences of representational drift. Current Opinion in Neurobiology, 58:141–147, Oct. 2019. ISSN 0959-4388. doi: 10.1016/j.conb.2019.08.005. URL https://www.sciencedirect.com/science/article/pii/S0959438819300303.

M. E. Rule, A. R. Loback, D. V. Raman, L. N. Driscoll, C. D. Harvey, and T. O’Leary. Stable task information from an unstable neural population. eLife, 9:e51121, July 2020. ISSN 2050-084X. doi: 10.7554/eLife.51121. URL https://elifesciences.org/articles/51121.

S. Sadeh and C. Clopath. Contribution of behavioural variability to representational drift. bioRxiv, 2022.

D. B. Salkoff, E. Zagha, E. McCarthy, and D. A. McCormick. Movement and performance explain widespread cortical activity in a visual detection task. Cerebral Cortex, 30(1):421–437, 2020.

C. E. Schoonover, S. N. Ohashi, R. Axel, and A. J. Fink. Representational drift in primary olfactory cortex. Nature, 594(7864):541–546, 2021a. ISSN 14764687. doi: 10.1038/s41586-021-03628-7. URL http://dx.doi.org/10.1038/s41586-021-03628-7.

C. E. Schoonover, S. N. Ohashi, R. Axel, and A. J. Fink. Representational drift in primary olfactory cortex. Nature, 594(7864):541–546, 2021b. ISSN 14764687. doi: 10.1038/s41586-021-03628-7. URL http://dx.doi.org/10.1038/s41586-021-03628-7. Publisher: Springer US.

M. G. Shuler and M. F. Bear. Reward timing in the primary visual cortex. Science, 311(5767): 1606–1609, 2006.

J. H. Siegle, X. Jia, S. Durand, S. Gale, C. Bennett, N. Graddis, G. Heller, T. K. Ramirez, H. Choi, J. A. Luviano, et al. A survey of spiking activity reveals a functional hierarchy of mouse corticothalamic visual areas. bioRxiv, page 805010, 2019.

J. H. Siegle, P. Ledochowitsch, X. Jia, D. J. Millman, G. K. Ocker, S. Caldejon, L. Casal, A. Cho, D. J. Denman, S. Durand, P. A. Groblewski, G. Heller, I. Kato, S. Kivikas, J. Lecoq, C. Nayan, K. Ngo, P. R. Nicovich, K. North, T. K. Ramirez, J. Swapp, X. Waughman, A. Williford, S. R. Olsen, C. Koch, M. A. Buice, and S. E. de Vries. Reconciling functional differences in populations of neurons recorded with two-photon imaging and electrophysiology. eLife, 10: 1–35, 2021. ISSN 2050084X. doi: 10.7554/eLife.69068.

K. K. Sit and M. J. Goard. Coregistration of heading to visual cues in retrosplenial cortex. preprint, Neuroscience, Mar. 2022. URL http://biorxiv.org/lookup/doi/10.1101/2022.03.25.485865.

C. Stringer, M. Pachitariu, N. Steinmetz, C. B. Reddy, M. Carandini, and K. D. Harris. Spontaneous behaviors drive multidimensional, brain-wide population activity. BioRxiv, page 306019, 2018.

J. Sugar, M. P. Witter, N. M. van Strien, and N. L. M. Cappaert. The retrosplenial cortex: intrinsic connectivity and connections with the (para)hippocampal region in the rat. An interactive connectome. Frontiers in neuroinformatics, 5:7, 2011. ISSN 1662-5196. doi: 10.3389/fninf.2011.00007. ISBN: 1662-5196 (Electronic)\n1662-5196 (Linking).

A. Thiele, A. Pooresmaeili, L. Delicato, J. Herrero, and P. Roelfsema. Additive effects of attention and stimulus contrast in primary visual cortex. Cerebral Cortex, 19(12):2970–2981, 2009. doi: 10.1093/cercor/bhp070.

S. True. Neural correlates of attention in primate visual cortex. Trends in Neurosciences, 24(5): 295–300, 2001. doi: 10.1016/s0166-2236(00)01814-2.

R. Vallat. Pingouin: statistics in python. Journal of Open Source Software, Feb. 2018. doi: https://doi.org/10.21105/joss.01026. URL https://pingouin-stats.org/.

T. Van Groen and J. M. Wyss. Connections of the retrosplenial granular b cortex in the rat. The Journal of comparative neurology, 463(3):249–63, Aug. 2003. ISSN 0021-9967. doi: 10.1002/cne.10757. URL http://www.ncbi.nlm.nih.gov/pubmed/12820159.

S. D. Vann, J. P. Aggleton, and E. A. Maguire. What does the retrosplenial cortex do? Nature Reviews Neuroscience, 10(11):792–802, 2009. ISSN 1471003X. doi: 10.1038/nrn2733.

Q. Wang, O. Sporns, and A. Burkhalter. Network analysis of corticocortical connections reveals ventral and dorsal processing streams in mouse visual cortex. The Journal of neuroscience: the official journal of the Society for Neuroscience, 32(13):4386–99, Mar. 2012. ISSN 1529-2401. doi: 10.1523/JNEUROSCI.6063-11.2012. URL http://www.pubmedcentral.nih.gov/articlerender.fcgi?artid=3328193&tool=pmcentrez&rendertype=abstract.

J. M. Wyss and T. Van Groen. Connections between the retrosplenial cortex and the hippocampal formation in the rat: a review. Hippocampus, 2(1):1–11, Jan. 1992. ISSN 1050-9631. doi: 10.1002/hipo.450020102. URL http://www.ncbi.nlm.nih.gov/pubmed/1308170.

